# A single-cell transcriptomics CRISPR-activation screen identifies new epigenetic regulators of zygotic genome activation

**DOI:** 10.1101/741371

**Authors:** Celia Alda-Catalinas, Danila Bredikhin, Irene Hernando-Herraez, Oana Kubinyecz, Fátima Santos, Mélanie A. Eckersley-Maslin, Oliver Stegle, Wolf Reik

## Abstract

Zygotic genome activation (ZGA) is a crucial developmental milestone that remains poorly understood. This first essential transcriptional event in embryonic development coincides with extensive epigenetic reprogramming processes and is orchestrated, in part, by the interplay of transcriptional and epigenetic regulators. Here, we developed a novel high-throughput screening method that combines pooled CRISPR-activation (CRISPRa) with single-cell transcriptomics to systematically probe candidate regulators of ZGA. We screened 230 epigenetic and transcriptional regulators by upregulating their expression with CRISPRa in mouse embryonic stem cells (ESCs). Through single-cell RNA-sequencing (scRNA-seq) of CRISPRa-perturbed cells, we generated approximately 200,000 single-cell transcriptomes, each transduced with a unique short-guide RNA (sgRNA) targeting a specific candidate gene promoter. Using integrative dimensionality reduction of the perturbation scRNA-seq profiles, we characterized molecular signatures of ZGA and uncovered 44 factors that promote a ZGA-like response in ESCs, both in the coding and non-coding transcriptome. Upon upregulation of these factors, including the DNA binding protein Dppa2, the chromatin remodeller Smarca5 and the transcription factor Patz1, ESCs adopt an early embryonic-like state. Supporting their roles as ZGA regulators, Dppa2 and Smarca5 knock-out ESCs lose expression of ZGA genes, however, overexpression of Dppa2 in Smarca5 knock-out ESCs, but not *vice versa*, rescues ZGA-like expression, suggesting that Smarca5 regulates ZGA upstream and via Dppa2. Together, our single-cell transcriptomic profiling of CRISPRa-perturbed cells provides comprehensive system-level insights into the molecular mechanisms that orchestrate ZGA.

**Highlights:**

- First large-scale screen combining pooled CRISPRa with scRNA-seq.
- Multi-omics factor analysis identifies a ZGA-like signature for 44 of the candidate regulators.
- Dppa2, Smarca5 and Patz1 were validated as strong inducers of ZGA gene expression.
- Smarca5 regulates zygotic genome activation in a Dppa2-dependent manner.

## Introduction

Zygotic genome activation (ZGA) is the first transcriptional event that takes place in an embryo (reviewed in Vastenhouw et al. 2019) and is a critical step in early development. In mouse, following an initial minor wave of ZGA in the late zygote, the major wave of ZGA occurs at the mid-to-late two-cell embryo stage and is characterized by the transcriptional activation of thousands of genes (reviewed in Vastenhouw et al. 2019; Jukam et al. 2017; Svoboda 2018.; Yartseva & Giraldez 2015). Significantly, in addition to the transcriptome, the epigenetic and chromatin landscape is drastically remodelled during this transition, including reprogramming of histone post-translational modifications, global chromatin accessibility and three-dimensional structure and global DNA demethylation (reviewed in Fraser & Lin 2016; Eckersley-Maslin, Alda-Catalinas & Reik 2018; Jansz & Torres-Padilla 2019). However, while several regulators of ZGA have been identified (reviewed in Eckersley-Maslin, Alda-Catalinas & Reik 2018), a comprehensive understanding of the complex regulation of the transcriptional and epigenetic events that occur during ZGA remains elusive.

High-throughput screening in preimplantation mouse embryos is not feasible due to the scarcity of material, maternal stores of proteins and complex manipulation techniques required. Recent studies have shown that a ZGA-like state can be mimicked in mouse ESCs (Macfarlan et al. 2012; Zalzman et al. 2010; Bošković et al. 2014; Ishiuchi et al. 2015; Akiyama et al. 2015; Eckersley-Maslin et al. 2016; Rodriguez-Terrones et al. 2018). Consequently, these cells represent an ideal system for *in vitro* screening and have been previously used to identify regulators of ZGA (Fu et al. 2019; Yan et al. 2019; Eckersley-Maslin et al. 2019; Rodriguez-Terrones et al. 2018). While most of these studies probing ZGA regulators in ESCs have focused on repressors (Fu et al. 2019; Rodriguez-Terrones et al. 2018), positive inducers of ZGA have thus far not been interrogated in a systematic manner. Such regulators are more relevant given the transcriptionally inactive state prior to ZGA and could be identified in ESCs by assessing the transcriptional changes triggered downstream of their overexpression (Eckersley-Maslin et al. 2019).

Recent studies have shown the power of combining pooled CRISPR/Cas9-based screening with single-cell RNA-sequencing (scRNA-seq) to obtain a comprehensive read-out, enabling interrogation of gene function and regulation at a cellular level in an unbiased manner (Jaitin et al. 2016; Dixit et al. 2016; Adamson et al. 2016; Datlinger et al. 2017; Xie et al. 2017; Genga et al. 2019; Gasperini et al. 2019; Replogle et al. 2018). However, these studies have all exclusively considered loss-of function perturbations through CRISPR knock-out (Jaitin et al. 2016; Dixit et al. 2016; Datlinger et al. 2017) or CRISPR-interference (CRISPRi) (Adamson et al. 2016; Xie et al. 2017; Genga et al. 2019; Gasperini et al. 2019; Replogle et al. 2018). Consequently, these existing approaches can only be used to interrogate genes that are already expressed in the cellular system under study (Gilbert et al. 2014).

CRISPR-activation (CRISPRa) is a potent tool for selective transcriptional upregulation of endogenous genes, which functions by targeting a dead Cas9 (dCas9) with transcriptional co-activators to gene promoters using short-guide RNAs (sgRNAs) (Cheng et al. 2013.; Gilbert et al. 2014; Chavez et al. 2015; Konermann et al. 2014), and has been successfully used for cellular reprogramming and the study of cellular transitions (Chakraborty et al. 2014; Black et al. 2016; Peng Liu et al. 2018; Weltner et al. 2018; Yang et al. 2019; Genga et al. 2019). CRISPRa is preferable to traditional overexpression techniques, such as cloned cDNA overexpression, as it leads to target gene activation at physiologically-relevant levels (Chavez et al. 2015; Sanson et al. 2018; Yang et al. 2019). Moreover, as it does not require cloning of genes, CRISPRa is highly scalable and allows the activation of genes that are, otherwise, difficult to clone or transfect into cells (Konermann et al. 2014; Joung et al. 2016; Horlbeck et al. 2016). The ability to combine CRISPRa screening with a transcriptome readout at single cell resolution would provide a novel high-throughput tool to comprehensively probe a large number of candidate genes to identify key regulators of transcriptional activation events, such as ZGA, while providing a comprehensive understanding of their function at the cellular level.

Here, we developed, for the first time, a high-throughput CRISPRa screening method that combines pooled sgRNA delivery with single-cell transcriptomic readout, and we applied this new technology to identify novel regulators of ZGA in ESCs. Using integrative dimensionality reduction analysis, we identified previously known and novel factors that induce ZGA-like transcription, including the binding protein Dppa2, the chromatin remodeller Smarca5 and the transcription factor Patz1. Furthermore, we mechanistically dissected a key part of the ZGA regulation network, revealing that Smarca5 regulates ZGA via Dppa2.

## Results

### A CRISPR-activation screen for ZGA regulators at single-cell resolution

To systematically identify novel regulators of ZGA, we developed a high-throughput pooled screening method that combines CRISPRa with single-cell transcriptomics (Fig. 1A). Given the large-scale epigenetic and transcriptional changes that occur during the maternal-to-zygotic transition (reviewed in Eckersley-Maslin, Alda-Catalinas & Reik 2018), we hypothesized that maternal epigenetic and transcriptional factors have crucial roles in regulating ZGA and consequently focused our screen on such candidate regulators. Due to the technical limitations of high-throughput screening in early embryos, we used mouse ESC as an *in vitro* proxy to mimic ZGA (Eckersley-Maslin, Alda-Catalinas & Reik 2018; Fu et al. 2019; Yan et al. 2019; Eckersley-Maslin et al. 2019; Rodriguez-Terrones et al. 2018), reasoning that overexpression of ZGA regulators may induce a ZGA-like transcriptional signature which can be captured by scRNA-seq.

**Figure 1:**
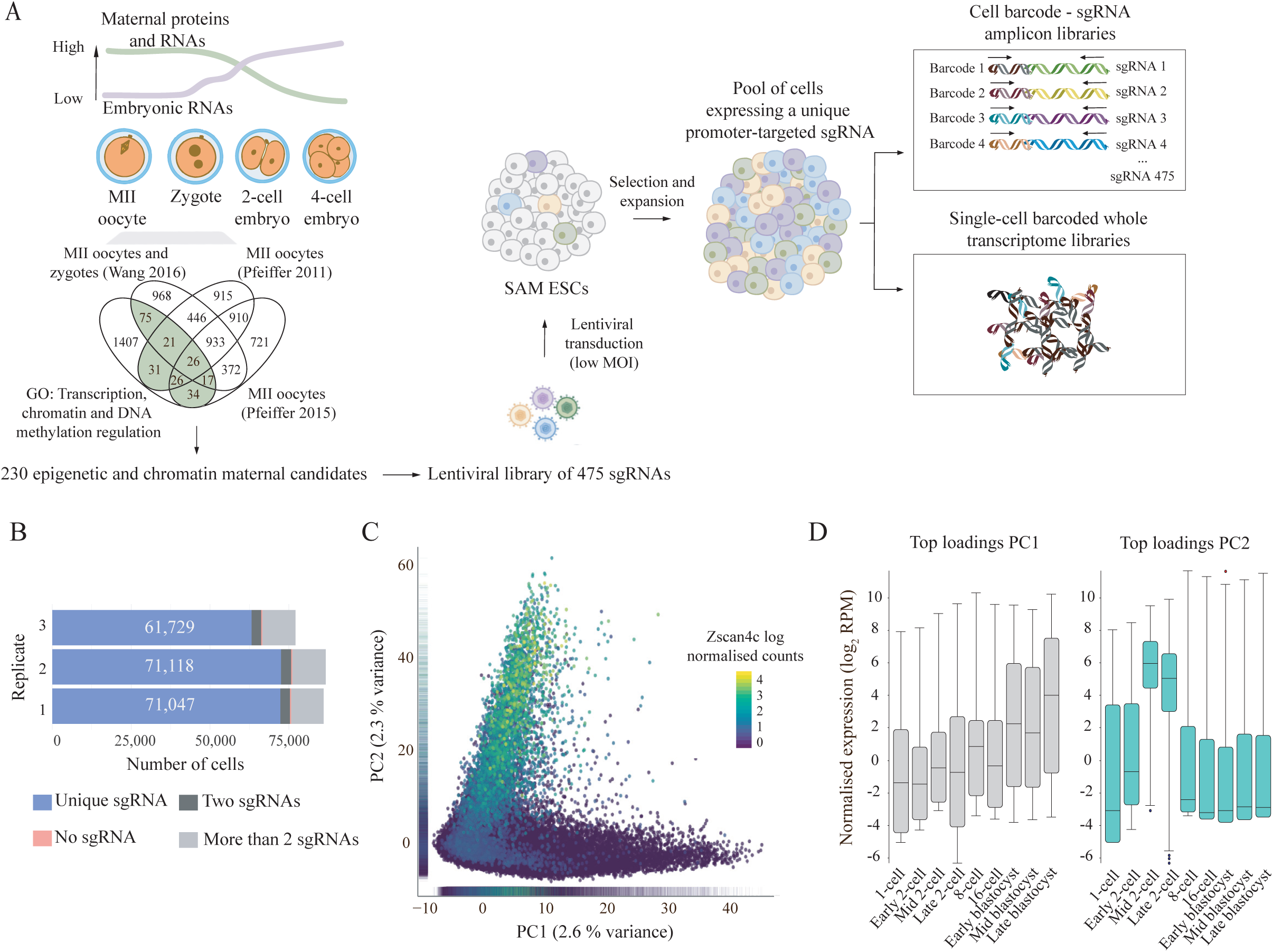
A CRISPR-activation screen for ZGA regulators at single-cell resolution. **A)** Schematic overview of the single-cell CRISPR-activation screen, highlighting the selection of candidates, lentiviral transduction strategy and generation of 10X Genomics 3’ single-cell RNA-sequencing libraries and barcoded sgRNA amplicon libraries. **B)** Number of cells expressing a unique sgRNA (blue), two sgRNAs (dark grey), more than two sgRNAs (light grey) or none (pink) in each of the three transduction replicates. The number of cells assigned to a unique sgRNA in each replicate is displayed. **C)** Principal component analysis of all cells that passed quality control, displaying a scatterplot of the first two principal components (PC1 versus PC2) with cells coloured by Zscan4c expression. Marginal distributions of PC1 and PC2 values are displayed as rug plots along the respective axis. **D)** Box-whisker plots showing normalised expression levels (log_2_ reads per million; RPM) for the top 50 loadings for PC1 (grey) and PC2 (light blue) during preimplantation development (data analysed from Deng et al. 2014) (see Table S3 for gene loadings). As expected in serum-grown ESCs, PC1 loadings peak at blastocyst stages whereas PC2 loadings peak at mid-to-late two-cell embryo stages, identifying this component as ZGA-like.

Our screening method builds on the robust and potent SAM CRISPRa system (Konermann et al. 2014) in which in addition to dCas9-VP64, high activation levels are achieved by recruiting the trans-activators p65 and HSF1, both fused to a MS2 RNA binding protein, through MS2 loops contained within the sgRNA scaffold sequence. Serum-grown clonal ESCs expressing both dCas9-VP64 and MS2-p65-HSF1 constitutively (referred to as SAM ESC) have a largely unchanged transcriptome compared to parental E14 ESCs (Fig. S1A), indicating that expression of the CRISPRa machinery does not substantially alter gene expression. To optimize and validate our system, we initially carried out a pilot experiment considering two strong candidate regulators, thereby confirming that CRISPRa can be used to induce a ZGA-like signature detectable by scRNA-seq. Briefly, SAM ESCs were transduced with sgRNAs targeting either the murine endogenous retrovirus with leucine tRNA primer (MERVL) long terminal repeats (LTRs) or zinc finger and SCAN domain containing 4 (Zscan4) gene cluster promoters (Table S1), and single cell transcriptomes were captured using the 10X Genomics scRNA-seq 3’ polyA-primed platform (Fig. S1B-D, see Materials and Methods). These key markers of ZGA are expressed in a low proportion of ESCs (Macfarlan et al. 2012; Zalzman et al. 2010; Eckersley-Maslin et al. 2016). CRISPRa significantly increased the proportion of cells expressing MERVL and Zscan4 3-fold (3.49% to 10.43%) and 2.8-fold (18.63% to 52.84%), respectively, compared to a non-targeting sgRNA control (Fig. S1E-F). Interestingly, MERVL LTR activation led to Zscan4 upregulation and *vice versa* (Fig. S1E-F), suggesting synergistic regulation as part of a network. We then defined a ZGA signature based on 2115 genes published in the literature to be expressed in the mouse embryo during ZGA or in the ZGA-like state of mouse ESCs (Eckersley-Maslin et al. 2016; Hendrickson et al. 2017; Y. Li et al. 2018) (Table S2). Notably, we found that the proportion of cells expressing these ZGA transcripts increased 4.28-fold and 3.34-fold upon MERVL LTR and Zscan4 CRISPRa, respectively (2% of cells transduced with a non-targeting sgRNA control to 8.56% of cells transduced with MERVL LTR sgRNAs and 6.67% of cells transduced with Zscan4 sgRNAs) (Fig. S1G). The upregulation of ZGA genes upon MERVL LTR activation is consistent with MERVL LTRs acting as functional promoters driving the expression of hundreds of chimeric ZGA transcripts (Macfarlan et al. 2012; Huang et al. 2017; Franke et al. 2017). Similarly, Zscan4c cDNA overexpression has recently been shown to induce the expression of ZGA genes (Eckersley-Maslin et al. 2019; Zhang et al. 2019). Collectively, the results from this pilot experiment not only validated 10X Genomics scRNA-seq as a suitable readout of ZGA-like expression following CRISPRa of relevant regulators, but also enabled us to estimate that approximately 400 cells per sgRNA (power 0.8, corrected p-value <0.0005, see Materials and Methods) are required to detect a ZGA-like transcriptional response upon CRISPRa of a positive hit exerting a similar effect to MERVL and Zscan4 activation.

Next, we applied our screening method to an extensive set of candidate regulators of ZGA shortlisted using publicly available proteomic datasets (Pfeiffer et al. 2011; Pfeiffer et al. 2015; Wang et al. 2016) and gene ontology enrichment. In total, we considered 230 proteins present in MII oocytes and zygotes with roles in transcription and epigenetic regulation (Fig. 1A and Table S1) which are expressed prior to and at the time of ZGA (Fig. S1H). Next, we designed a pooled sgRNA library containing two sgRNAs for each of the 230 candidate maternal ZGA regulators, targeting the 180 base-pair (bp) window upstream of the transcription start site (TSS), along with fifteen non-targeting sgRNA controls (Konermann et al. 2014; Joung et al. 2016) (Table S1). The resulting library consisting of 475 sgRNAs was cloned into a lentiviral vector modified from CROP-seq (Datlinger et al. 2017) to include MS2 loops in the sgRNA scaffold sequence (referred to as CROP-sgRNA-MS2, Fig S1I, see Materials and Methods). This lentiviral vector backbone enables both CRISPRa via SAM and capture of the sgRNA target sequence in 10X Genomics 3’ scRNA-seq libraries. Notably, the cloned plasmid library showed representation of the 475 sgRNAs (Fig. S1J, Table S1).

SAM ESCs were transduced with this lentiviral library of 475 sgRNAs at a <0.1 multiplicity-of-infection (MOI) in triplicate (Fig. S2A). Following selection and expansion of the pool of transduced cells, single-cell transcriptomes were generated using 10X Genomics scRNA-seq 3’ polyA-primed platform and their corresponding sgRNAs were further amplified using a specific amplification protocol (Hill et al. 2018) (Fig. 1A, see Materials and Methods). A total of 341,103 single-cell transcriptomes were sequenced across the three transduction replicates (see Materials and Methods). After scRNA-seq quality controls (Fig. S2B-2D), sgRNA assignment to each individual cell and removal of cells with no or multiple assigned sgRNAs, we obtained a total of 203,894 cells expressing a unique sgRNA for further analysis (Fig. 1B, see Materials and Methods). All sgRNAs were captured consistently across the three replicates, with an average coverage of 437 cells per sgRNA for the combined dataset (Fig. S2E-F). The number of cells expressing each sgRNA (Table S1) matched the representation distribution of the sgRNA plasmid library (Fig. S2G), indicating that activation of the target genes did not have any strong effects on cell proliferation or viability.

Initially, we performed principal component analysis (PCA) to explore the major sources of variation in our dataset (Fig. 1C). While gene ontology enrichment of the top 50 gene loadings of the first principal component (PC1) identified this component as capturing intrinsic variation in cell-to-cell contacts and cell shape (Fig. S2H, Table S3), excitingly, the second component (PC2) robustly captured variation of genes that are highly expressed in mid-to-late two-cell embryos at the time of ZGA, including Zscan4c, Zscan4d, Gm8300 and Tmem92 (Fig. 1C-D, S2H-I, Table S3). This was consistent between replicates (Fig. S2J), validating the robustness of our screen. Together, these results show that our CRISPRa scRNA-seq screen induced expression variation that mimics a ZGA-like transcriptional response in ESCs, suggesting that a substantial fraction of our selected maternal candidates did indeed induce a ZGA-like gene signature.

### Identification of activators of a ZGA-like transcriptional signature

Next, we set to characterize the observed ZGA-like transcriptional signature in more detail. In addition to coding genes, we also included transposable or repeat elements in our analysis (Fig. S3A-B, see Materials and Methods), since they are key drivers of gene expression during early embryonic development (reviewed in Rodriguez-Terrones & Torres-Padilla 2018). We used multi-omics factor analysis (MOFA) (Argelaguet et al. 2018) to combine the expression of coding genes and repeat elements within a single model, and to disentangle individual activating sgRNAs responsible for inducing the observed ZGA-like response (see Materials and Methods). Briefly, the MOFA framework allows integration of both data modalities as distinct views and it identifies the most important factors that explain the transcriptional variability within the dataset (Fig. 2A, see Materials and Methods). Excitingly, amongst the MOFA factors identified (Table S4), factor 3 again captured a ZGA-like signature: the coding genes that have the highest loadings for factor 3 are ZGA genes highly expressed in mid-to-late two-cell embryos (Fig. 2B-C, S3C-F) and the ZGA-related MERVL repeat (Macfarlan et al. 2012) was most prominently associated to factor 3 amongst the repeat classes analysed (Fig. 2D, S3G).

**Figure 2:**
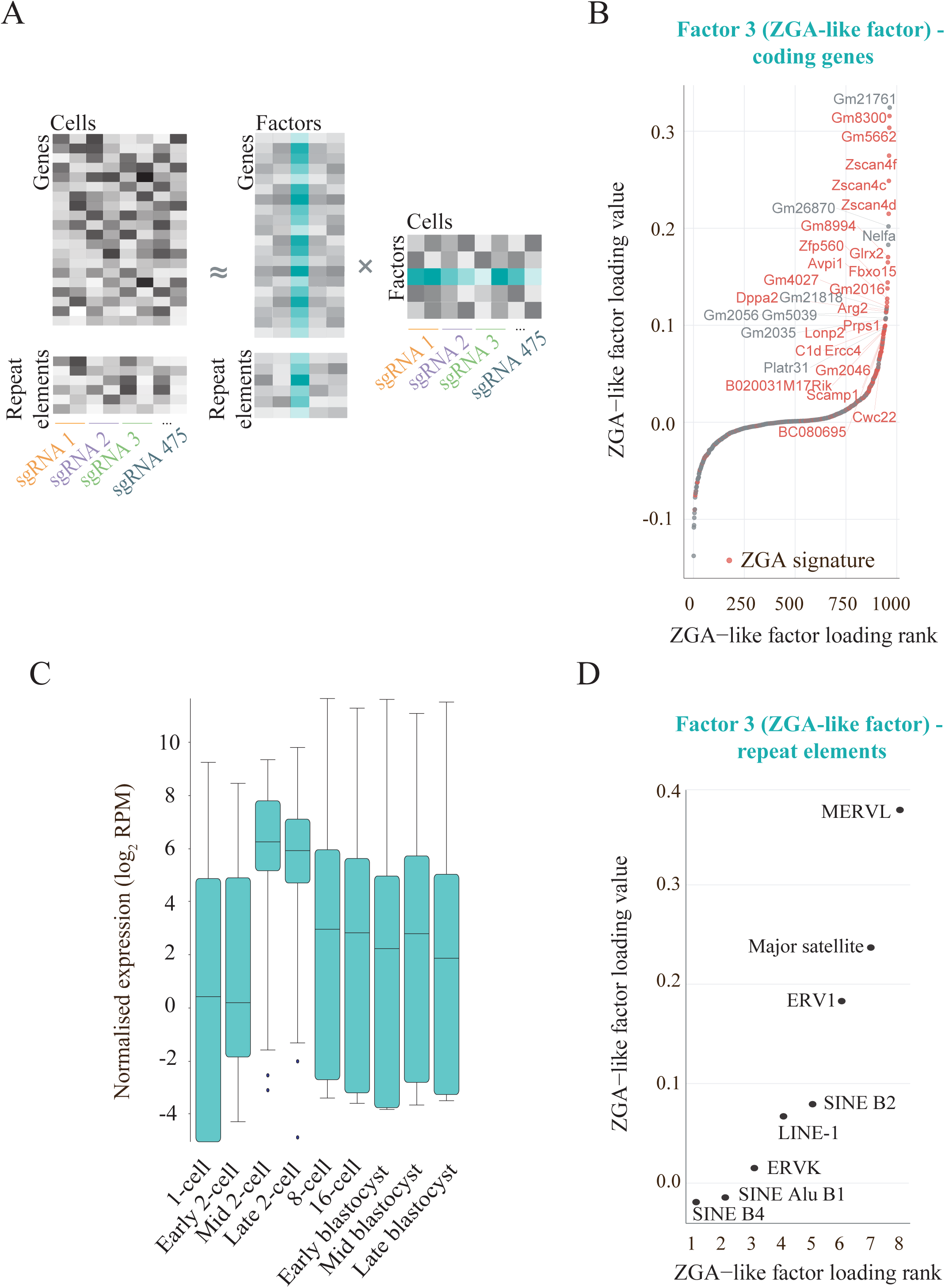
Identification a ZGA-like transcriptional signature using MOFA. **A)** Schematic of the joint analysis of coding gene and repeat elements expression using multi-omics factor analysis (MOFA). Data matrices of dimension features (genes or repeat elements) in cells grouped by sgRNA expression are treated as distinct views in the model and decomposed into the product of weights (or loadings) and factors. Factor 3 in the trained model, interpreted as a ZGA-like factor, is highlighted in green. **B)** Coding genes ranked by their loadings of MOFA factor 3, highlighting in red previously known ZGA genes (as described in Table S2, see also Table S4 for gene loading values), thus identifying this factor as ZGA-like. **C)** Box-whisker plots showing normalised expression levels (log_2_ reads per million; RPM) for the top 50 gene loadings of MOFA factor 3 (ZGA-like factor) during preimplantation development (data analysed from Deng et al. 2014) (see Table S4 for gene loadings). **D)** Repeat element families ranked by their loadings of MOFA factor 3 (ZGA-like factor).

To reveal individual candidate genes that induced a ZGA-like signature when activated, we grouped the cells by expression of the targeting sgRNA and ranked each group based on its contribution to the overall transcriptional variance explained by the MOFA ZGA-like factor or factor 3 (Table S1, see Materials and Methods). This identified 46 sgRNAs targeting 44 unique genes for which the MOFA factor explained a larger fraction of variance than for the fifteen non-targeting sgRNAs controls (Fig. 3A, Table S1, see Materials and Methods). Importantly, cells expressing these 46 sgRNAs specifically induced the expression of ZGA genes associated to factor 3 (Fig. 3B, S4A), while genes associated to other MOFA factors remained largely unaltered (Fig. S4B), highlighting the specificity of these screen hits in upregulating a ZGA-like signature. Similarly, the ZGA-associated MERVL elements (Macfarlan et al. 2012), major satellite repeats (Casanova et al. 2013), ERV1 (Zhang et al. 2019) and SINE B2 elements (Vasseur et al. 1985), but not other repeat families, were upregulated by these sgRNA hits (Fig. 3C, S4C), consistent with these repeat families ranking top in the MOFA ZGA-like factor (Fig. 2D).

**Figure 3:**
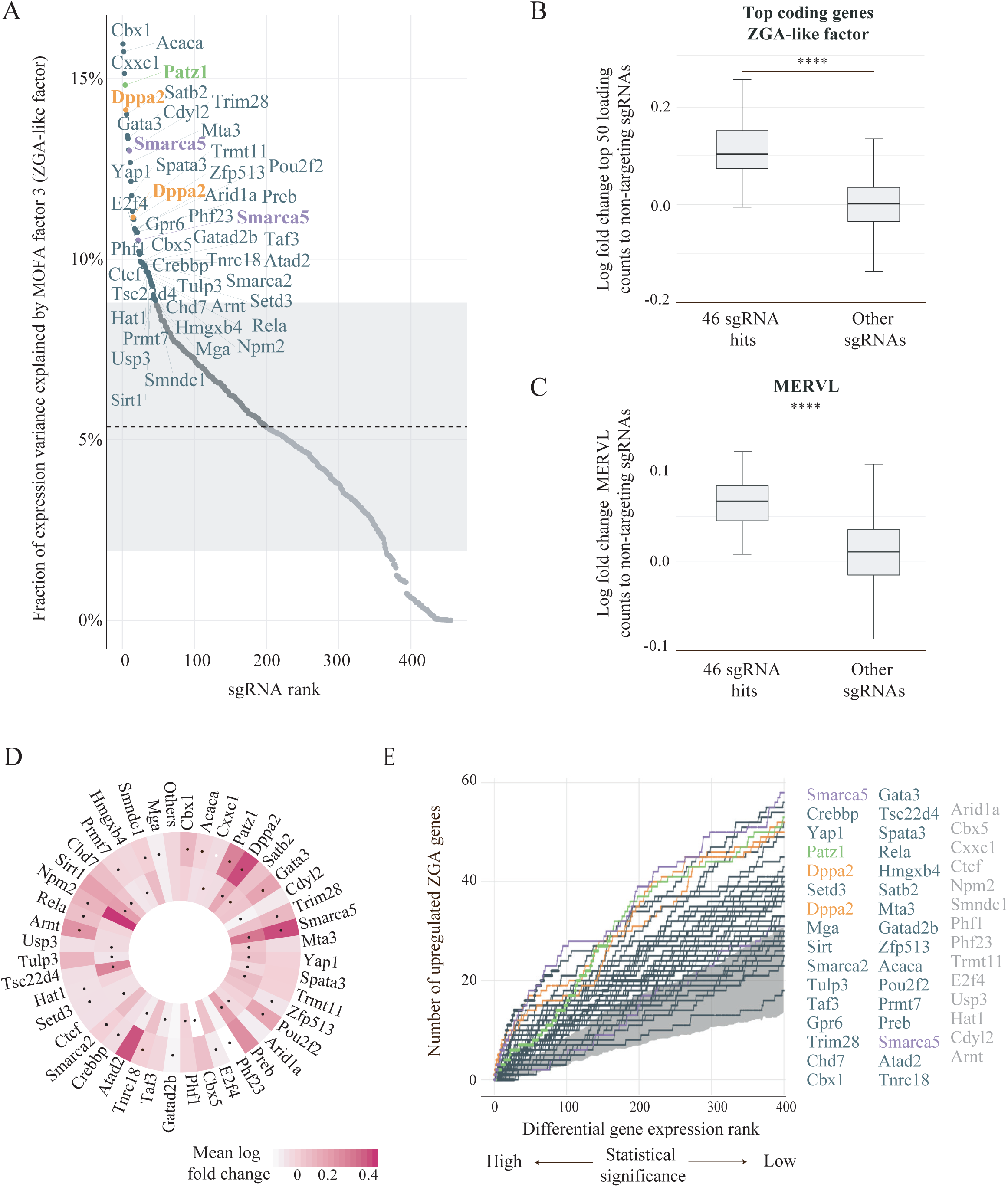
Identification of activators of a ZGA-like transcriptional signature. **A)** Ranking of the 460 candidate sgRNAs by the fraction of expression variance (average of coding genes and repeat elements) explained by the MOFA factor 3 (ZGA-like factor). The fraction of expression variance explained by the MOFA factor 3 for the fifteen non-targeting sgRNA controls is depicted in the background as mean (dashed line) and plus and minus one standard deviation (shaded area). The names of the target genes for sgRNAs that exceed the variance explained by non-targeting sgRNA controls are shown (see Table S1 for the full ranking) with Dppa2 (orange), Smarca5 (purple) and Patz1 (green) sgRNA hits highlighted. **B)** Box-whisker plots showing log fold change expression for the top 50 genes ranked by the MOFA factor 3 (ZGA-like factor) loadings in cells expressing the 46 sgRNA hits and cells expressing other targeting sgRNAs, compared to cells expressing non-targeting sgRNA controls. Expression is quantified in normalised counts (****: p-value =10^-22^, Mann-Whitney two-sided test). **C)** Box-whisker plots showing log fold change of MERVL normalised counts in cells expressing the 46 sgRNA hits and cells expressing other targeting sgRNAs, compared to cells expressing non-targeting sgRNA controls (****: p-value =10^-16^, Mann-Whitney two-sided test). **D)** Target gene activation for the 46 sgRNA hits, measured by log fold change expression between cells expressing the relevant targeting sgRNA and cells expressing non-targeting sgRNA controls; both sgRNAs targeting the gene of interest are shown (outer and inner circle) and sgRNAs that were identified as hit for inducing a ZGA-like signature are highlighted with a black dot. **E)** Cumulative rank of the number of ZGA signature genes (as described in Table S2) upregulated by each sgRNA hit compared to non-targeting sgRNA controls, considering the top 400 genes ranked by statistical significance of differential gene expression test (generalised linear model likelihood ratio test as implemented in EdgeR, FDR<1). The background distribution based on differential gene expression between cells with non-targeting sgRNA controls is shown in grey, displaying plus and minus one standard deviation around the mean of ZGA signature genes recovered by non-targeting sgRNAs. The names of the target genes for sgRNAs identified as hits in **A)** are depicted, with those for which the differential gene expression rank overlaps with the non-targeting control background shown in grey. Dppa2 (orange), Smarca5 (purple) and Patz1 (green) sgRNA hits are highlighted.

Lastly, we investigated individual target genes induced by CRISPRa of the 46 sgRNA hits compared to non-targeting sgRNA controls (see Materials and Methods). We detected gene activation of the corresponding target gene for 26 of these 46 sgRNAs (56.5%, mean log fold change to non-targeting sgRNA controls >0) (Fig. 3D, Table S1), where lack of target gene activation for the remaining sgRNAs could be due to technical drop-outs that are commonly observed in scRNA-seq data. When assessing differential gene expression between targeted and non-targeted cells transcriptome-wide, only a small subset of genes was significantly differentially expressed for most hits (between 0 and 224, median 4.5; FDR<0.1, Table S1, see Materials and Methods). However, ranking of the top 400 upregulated genes by statistical significance identified the downstream genes of 32 out of the 46 sgRNA hits (69.6%) as being prominently enriched for known ZGA transcripts, compared the background level of enrichment calculated by differential gene expression between cells with non-targeting sgRNA controls (Fig. 3E, Tables S1 and S2). This result confirms the MOFA analysis, highlights its advantages in capturing relevant gene signatures, of both coding and non-coding transcription, in an unbiased way, and shows that MOFA can identify screen hits that are otherwise missed using differential gene expression ranking, due to lack of power to detect the effects on individual genes in a pre-defined signature. Interestingly, both sgRNAs targeting the DNA binding protein Dppa2 and the ATPase subunit of the ISWI chromatin remodeller complex Smarca5 (also known as Snf2h) induced a ZGA-like response (Fig. 3A, 3E). In summary, using MOFA to integrate the expression of coding genes and transposable elements in our CRISPRa scRNA-seq dataset, we identified 44 genes whose activation induces a ZGA-like transcriptional response. These include four previously known maternal ZGA regulators, namely Dppa2 (Eckersley-Maslin et al. 2019; De Iaco et al. 2019; Yan et al. 2019), Gata3 (Eckersley-Maslin et al. 2019), Yap1 (Yu et al. 2016) and Ctcf (Wan et al. 2008), and excitingly, 40 novel maternal proteins not previously linked to ZGA.

### Dppa2, Smarca5 and Patz1 regulate expression of ZGA transcripts

Next, we used two experimental approaches to validate Dppa2 and Smarca5, for which both targeting sgRNAs were hits, and the top-ranking transcription factor Patz1 (Fig. 3A, 3E, S5A). We also included Carhsp1 as a negative control, since both of its targeting sgRNAs showed effective target gene activation (Fig. S5B, Table S1) and ranked within the non-targeting sgRNA control background (Fig. S5A, Table S1). In these validation experiments, target gene overexpression was induced either by individual CRISPRa with one of the sgRNAs used in the screen (Table S1), or by cDNA-eGFP overexpression, and the transcriptomic changes were assessed by bulk polyA-capture RNA-sequencing in biological triplicates (Fig. 4A). Both methods led to target gene upregulation, similar to what was observed by scRNA-seq (Fig. S5B, S5C), and we observed markedly similar genome-wide transcriptional response patterns across methods (Fig. S5D). Importantly, we found that, despite the increased power in calling differential gene expression using bulk RNA-sequencing, the transcriptional changes captured by scRNA-seq and bulk RNA-sequencing upon CRISPRa of these four targets were highly correlated (Fig. S5E), demonstrating the robustness of our method. Together, these validation experiments revealed that CRISPRa with scRNA-seq readout is a robust method to assess the transcriptional responses triggered by gene overexpression.

**Figure 4:**
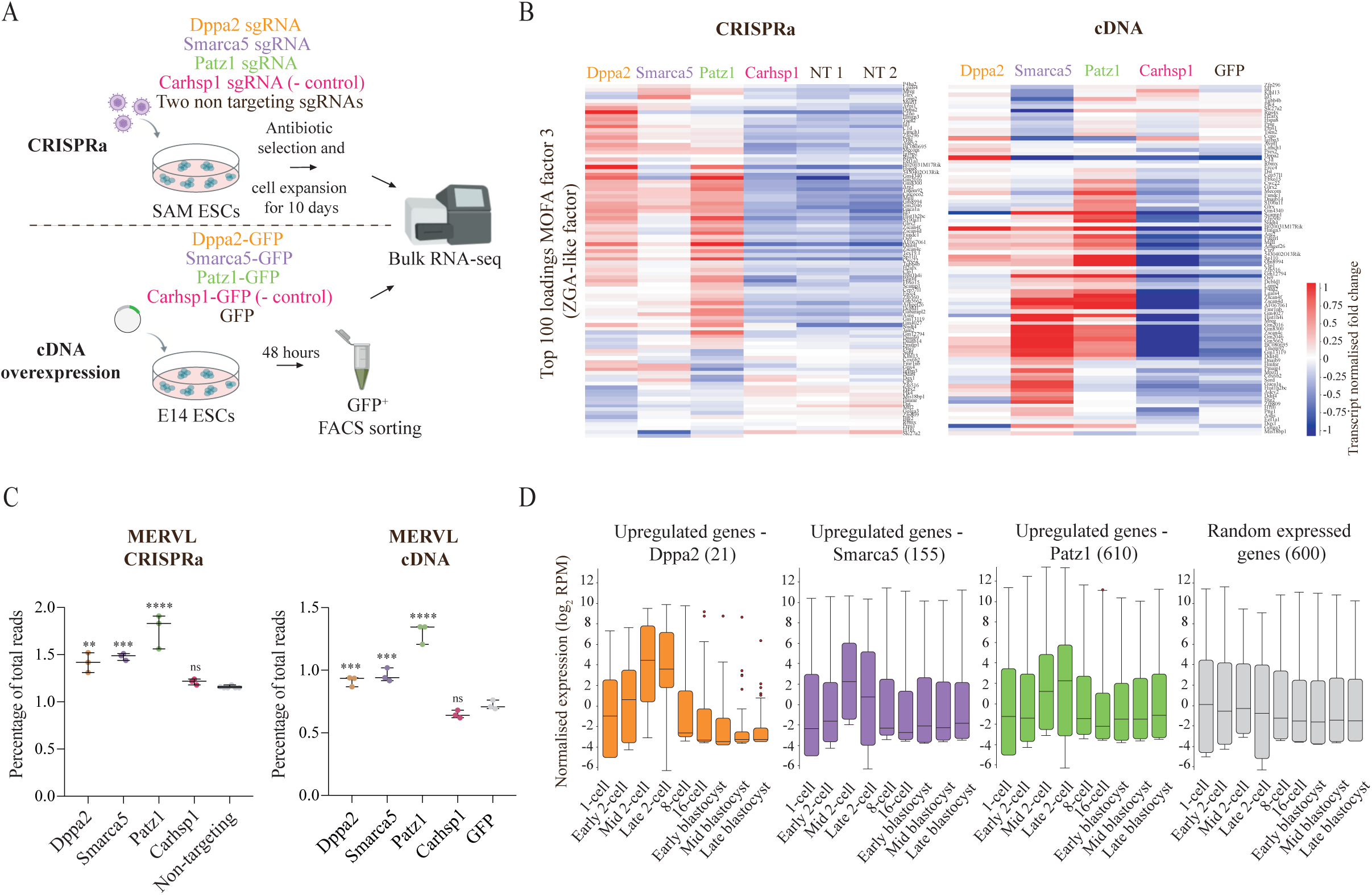
Validation of Dppa2, Smarca5 and Patz1 as ZGA regulators. **A)** Schematic representation of two sets of validation approaches to confirm the screen hits Dppa2, Smarca5 and Patz1, using Carhsp1 as a negative control hit, by CRISPRa (top) and cDNA overexpression (bottom) followed by bulk polyA RNA-sequencing. The sgRNAs used for CRISPRa in these bulk RNA-sequencing experiments are described in Table S1. **B)** Heatmap showing normalised gene expression, scaled per gene, of the top 100 loading genes for MOFA factor 3 (ZGA-like factor), in bulk RNA-sequencing libraries for Dppa2, Smarca5, Patz1, Carhsp1 CRISPRa (left) and cDNA overexpressing cells (right). Controls are two different non-targeting sgRNAs (NT1 and NT2) for CRISPRa and GFP only overexpression for cDNA. **C)** Box-whisker plots showing expression of the MERVL repeat family in percentage of total reads measured by bulk RNA-sequencing after CRISPRa (left) and cDNA overexpression (right) of Dppa2 (orange), Smarca5 (purple), Patz1 (green) and Carhsp1 (pink) and in controls (two non-targeting sgRNA controls for CRISPRa (left) and GFP^+^ only control for cDNA overexpression (right)). Each dot represents a biological replicate. Statistically significant differences to controls are reported as ****: p-value < 0.0001, ***: p-value <0.001, **: p-value <0.01, ns: non-significant; homoscedastic two-tailed t-test. **D)** Box-whisker plots showing normalised expression levels (log_2_ reads per million; RPM) of differentially upregulated genes by both CRISPRa and cDNA overexpression of Dppa2 (orange), Smarca5 (purple), Patz1 (green), and Carhsp1 (pink), as well as a random set of expressed genes (grey) during preimplantation development (data analysed from (Deng et al. 2014). Differential gene expression was calculated with EdgeR (FDR <0.5).

CRISPRa or cDNA overexpression of Dppa2, Smarca5 and Patz1, but not Carhsp1, led to upregulation of the genes captured in MOFA ZGA-like factor (Fig. 4B). Additionally, whole transcriptome-wide, these three hits triggered a strikingly similar transcriptional response which was clearly distinct from the changes triggered by Carhsp1 (Fig. S5D), suggesting they regulate similar transcriptional networks. Furthermore, MERVL, which captured the highest variability amongst the repeat families in the scRNA-seq dataset as analysed by MOFA (Fig. 2D), was significantly upregulated by Dppa2, Smarca5 and Patz1, but not by Carhsp1 (Fig. 4C). Interestingly, LINE-1 expression, which has also been linked to ZGA (Percharde et al. 2018), was induced by Dppa2 and Patz1 (Fig. S5F). Notably, genes significantly upregulated by both CRISPRa and cDNA overexpression of Dppa2, Smarca5 and Patz1 (see Materials and Methods) were highly expressed at the time of ZGA *in vivo* (Fig. 4D), providing strong evidence that these maternal factors activate a ZGA-like programme in ESCs. In summary, we confirmed three top screen hits as true regulators of ZGA-like transcription, validating our novel method as a reliable high-throughput tool for detecting positive regulators of transcriptional programmes.

### Smarca5 induces ZGA-like expression via Dppa2

Lastly, we focused on the mechanistic regulation of the two top hits Dppa2 and Smarca5. Dppa2 has recently been identified as a potent inducer of ZGA networks (Eckersley-Maslin et al. 2019; De Iaco et al. 2019; Yan et al. 2019). However, Dppa2 is also expressed in cells that do not express ZGA transcripts, suggesting additional chromatin remodellers are required for Dppa2’s ZGA-like function (Eckersley-Maslin et al. 2019). Smarca5 knock-down in zygotes reduces transcription of a set of genes (Torres-Padilla & Zernicka-Goetz 2006), but its role as a ZGA regulator has not been characterized yet. Both Dppa2 and Smarca5 are expressed throughout preimplantation development, with Smarca5 being more highly expressed in the oocyte than Dppa2, which increases in its expression from the two-cell stage (Fig. 5A). Strikingly, in zygotes, while Dppa2 localizes in the cytoplasm, Smarca5 is present in both pronuclei (Fig. 5B). However, in two-cell embryos, at the time of ZGA, Dppa2 translocates to the nucleus and both proteins co-localize (Fig. 5B-C). Importantly, Dppa2 and Smarca5 proteins have been shown to physically interact in ESCs (Hernandez et al. 2018), suggesting they may function together to regulate their ZGA target genes. Consistent with our overexpression results, analysis of recently published Smarca5 KO transcriptome data (Barisic et al. 2019) revealed that loss of Smarca5 led to downregulation of ZGA transcripts (Fig. 5D). Similarly, recent studies have also shown that Dppa2 KO ESCs lack expression of ZGA-like genes (Eckersley-Maslin et al. 2019; De Iaco et al. 2019; Yan et al. 2019). We next investigated whether Smarca5 exerts its ZGA regulatory function through its catalytic ATPase activity or through interactions with accessory subunits of the ISWI complex. Analysis of published RNA-sequencing data of Smarca5 KO ESCs (Barisic et al. 2019) revealed that wild-type, but not a catalytically-dead Smarca5 mutant, was able to restore expression of the 391 ZGA genes downregulated upon Smarca5 loss (Fig. 5D). This indicates that the regulation of ZGA by Smarca5 is dependent on its ATPase activity.

**Figure 5:**
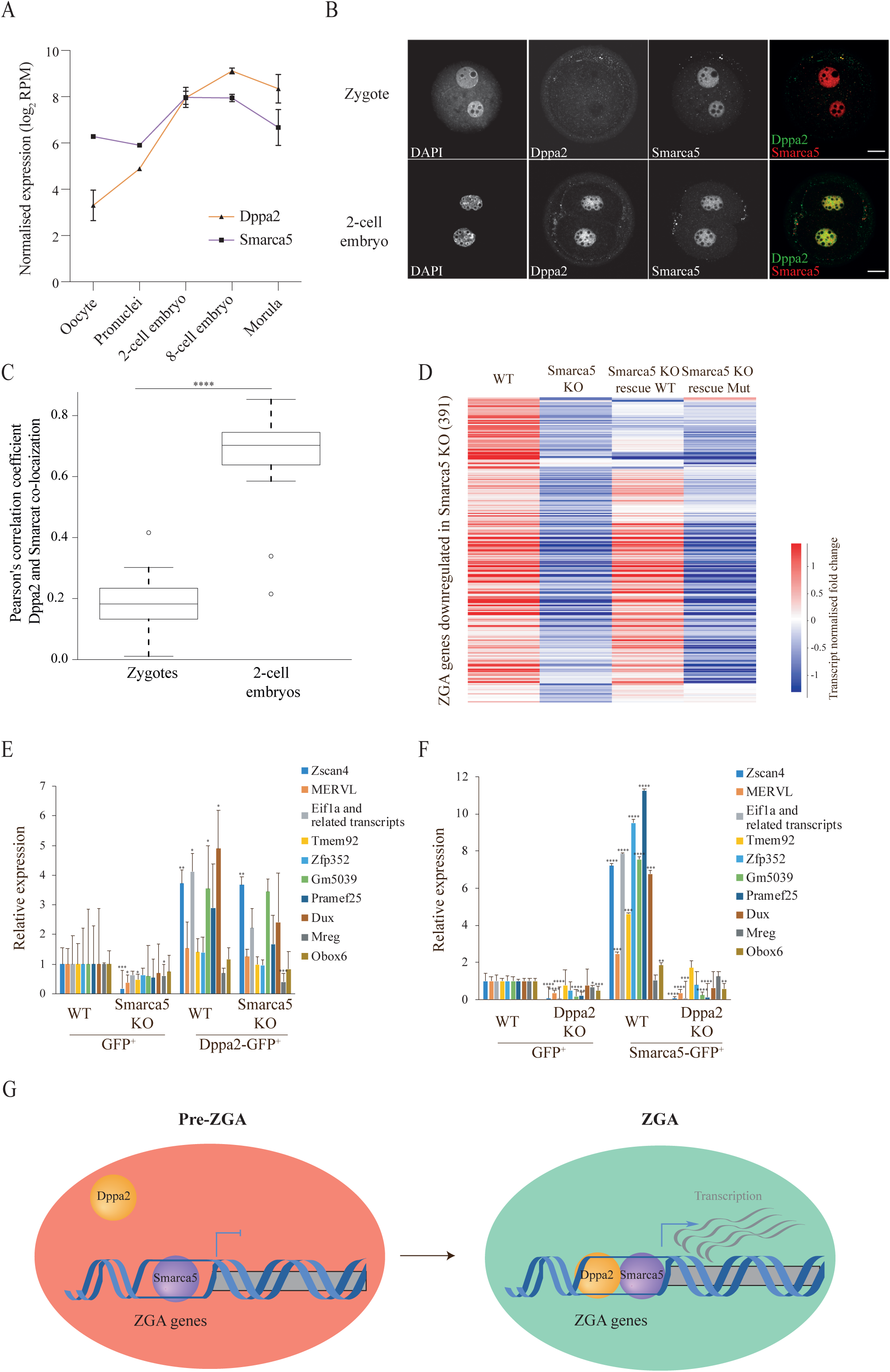
Smarca5 induces ZGA-like expression via Dppa2. **A)** Normalised expression levels (log_2_ reads per million; RPM) of Dppa2 (orange triangles) and Smarca5 (purple squares) in oocytes and preimplantation development (data analysed from Xue et al. 2013). Data is shown as mean plus standard deviation of biological replicates. **B)** Representative single optical slices of zygotes (top row) and 2-cell stage embryos (bottom row) immunostained for Dppa2 (green) and Smarca5 (red). Scale bars represent 20 µm. **C)** Box-plots showing Pearson correlation coefficients calculated for co-localization of Dppa2 and Smarca5 in the pronuclei of 10 zygotes and in the nuclei of 10 2-cell stage embryos. Co-localization values in the two pronuclei in zygotes and nuclei of each blastomere in two-cell embryos were measured separately. Dppa2 and Smarca5 co-localize in two-cell embryos but not in zygotes (****: p-value < 0.0001, Mann-Whitney test). **D)** Heatmap showing normalised expression, scaled per gene, of downregulated ZGA genes in Smarca5 mouse knock-out (KO) cells compared to wild-type (WT) (EdgeR, FDR<0.5), in WT ESCs, Smarca5 KO ESCs and Smarca5 KO ESCs expressing a Smarca5 WT protein or a Smarca5 catalytic-dead mutant protein (Mut) (data analysed from Barisic et al. 2019). **E), F)** Analysis of ZGA-like transcripts relative expression levels by quantitative reverse transcription PCR in **E)** WT and Smarca5 KO mouse ESCs after 48 hours transient transfection of GFP or Dppa2-GFP and **F)** WT and Dppa2 KO mouse ESCs after 48 hours transient transfection of GFP or Smarca5-GFP. GFP^+^ cells were FACS-sorted before gene expression analysis. Relative expression levels are normalised to WT cells transfected with GFP and sorted for GFP^+^. Data is shown as mean plus standard deviation of three biological replicates. Statistically significant differences to WT GFP^+^ control are reported (homoscedastic two-tailed t-test, **: p-value <0.01, ***: p-value <0.001, ****: p-value < 0.0001). **G)** Model of Dppa2 and Smarca5 action in regulation of ZGA: in a pre-ZGA state, the chromatin of ZGA genes is bound by Smarca5 (purple) and transcription is inactive; ZGA is triggered when Dppa2 is recruited to ZGA genes, inducing transcription that is facilitated by an open chromatin state induced by Smarca5.

Finally, given their co-localization in two-cell embryos (Fig. 5B-C), physical interaction in ESCs (Hernandez et al. 2018) and similar transcriptional effects (Fig. 4B-D, S5D), we sought to understand the mechanistic interdependencies, if any, between Dppa2 and Smarca5 through a series of KO and overexpression experiments analysed by quantitative reverse transcription PCR. First, to test whether Smarca5 is required for Dppa2’s function, we overexpressed Dppa2-eGFP in wild-type (WT) and Smarca5 KO cells (Fig. S6A-B). As shown by RNA-sequencing (Fig. 5D), expression of ZGA genes was downregulated in Smarca5 KO cells, although not completely lost (Fig. 5E). Excitingly, expression of these genes was partially rescued by Dppa2 overexpression (Fig. 5E), while the pluripotency gene Oct4 remained unaltered (Fig. S6C). This suggests that Smarca5 may act upstream or in parallel to Dppa2. To test between these options, we overexpressed Smarca5-eGFP in WT and Dppa2 KO ESCs (Fig. S6D-E). As expected, in WT cells, Smarca5 overexpression strongly induced the expression of ZGA genes, including the Zscan4 cluster and MERVL (Fig. 5F, S6F), while the pluripotency gene Oct4 remained unchanged (Fig. S6G). Interestingly, in Dppa2 KO ESCs, which have nearly absent levels of ZGA transcripts (Eckersley-Maslin et al. 2019) (Fig. 5F), ZGA-like gene expression could not be induced by Smarca5-eGFP overexpression (Fig. 5F). This suggests Dppa2 is required for Smarca5-mediated regulation of ZGA. Therefore, while Smarca5 requires Dppa2 for its ZGA transcriptional effects, Dppa2 does not require Smarca5. Notably, Smarca5 induces Dppa2 expression but not vice versa (Fig S6B, S6E). This is likely via a transcriptional feedback, loop as Dppa2 is also zygotically expressed during major ZGA (Fig. 5A, Table S2). In conclusion, we propose a model in which Dppa2 and Smarca5 act together to induce ZGA via chromatin remodelling and activation of transcription (Fig. 5G). Before ZGA, Smarca5 is already present in the nucleus, maintaining a poised open chromatin state of the promoters of ZGA genes; upon translocation of Dppa2 to the nucleus in the two-cell embryo, Smarca5 facilitates Dppa2 binding and activation of ZGA genes (Fig. 5G).

## Discussion

Here, we developed a novel pooled CRISPRa screen coupled with scRNA-seq readout, and applied this tool to identify new regulators of ZGA. Our dataset, comprised of 203,894 single ESCs each expressing a unique sgRNA, allowed us to investigate the transcriptional consequences following CRISPRa of 230 maternally-expressed epigenetic and transcriptional factors (Fig. 1A-B). By performing integrative dimensionality reduction of expression of both coding mRNAs and repeat elements (Fig. 2A), we revealed 44 maternal factors that induced a transcriptional response reminiscent of the major wave of ZGA (Fig. 3A-C). Dppa2, Smarca5 and Patz1 were validated, using complementary experimental approaches, as inductors of ZGA-like transcription in ESCs (Fig. 4B-D). These data not only validate our screen approach but also the roles of these proteins in regulating ZGA expression. Finally, we disentangled the interdependencies between Dppa2 and Smarca5 and put forth a model by which Smarca5 opens up the chromatin and facilitates ZGA via Dppa2 (Fig. 5G).

Our novel screening method provides a powerful way to systematically interrogate a large number of genes for their effects on specific transcriptional programmes, and we anticipate that it will be widely adaptable to many other biological contexts and research questions. While overexpression screens using traditional open reading frame (ORF) libraries have already been coupled with scRNA-seq readout (Parekh et al. 2018), CRISPRa has key advantages over traditional cDNA overexpression. Firstly, target genes are upregulated at physiologically-relevant levels (Chavez et al. 2015; Sanson et al. 2018; Yang et al. 2019). Secondly, it allows activation of genes that might be otherwise difficult to clone, as well as other genomic features such as repeat elements (Fig. S1E). Strikingly, despite the differences in gene dosage (Fig. S5C) and experimental design (Fig. 4A) in our validation experiments with CRISPRa and cDNA overexpression, the transcriptional changes triggered by our top hits Dppa2, Smarca5 and Patz1 are remarkably similar in the induction of ZGA gene expression (Fig. 4B-D, S5D), confirming CRISPRa as a robust method to analyse ZGA-like transcription.

Screening using single-cell transcriptomics has substantial advantages in terms of scalability and the possibility to disentangle cell-to-cell heterogeneity. However, it comes at a cost in terms of sensitivity for detecting transcriptional changes in individual cells (reviewed Kelsey et al. 2017). Related work using CRISPR KO CROP-seq (Datlinger et al. 2017) applied computational downsampling analysis to show that only 12-13 cells per sgRNA can suffice to detect the expected transcriptional signatures upon deletion of T-cell receptor signaling regulators. In our CRISPRa screen for ZGA regulators, we performed *a priori* power calculations and estimated that at least 400 cells were required per sgRNA to confidently detect a ZGA-like transcriptional response (Fig. S1G, Materials and Methods). While these design choices are highly dependent on the biological context of investigation, it is crucial to determine these parameters using prior knowledge or pilot data, as done in our study. Furthermore, to mitigate the reduced sensitivity of scRNA-seq, we considered a ZGA transcriptional signature rather than quantification of individual genes. To this end, our analysis builds on the MOFA framework (Argelaguet et al. 2018) to incorporate both the coding transcriptome and repetitive elements to define a robust and sensitive signature of ZGA-like transcriptional responses following CRISPRa (Fig. 2A). We demonstrate that this approach is superior in identifying relevant screen hits, compared to conventional differential expression analysis (Fig. 3A, 3E). Specific repeat classes are expressed during ZGA in mouse embryos, including MERVL (Kigami 2002; Peaston et al. 2004; Macfarlan et al. 2012) and LINE-1 retrotransposons (Percharde et al. 2018; Jachowicz et al. 2017). Consistently, our top hits Dppa2, Smarca5 and Patz1 all showed consistent upregulation of MERVL in the scRNA-seq and validation experiments, and Dppa2 and Patz1 also upregulated LINE-1 expression (Fig. 3C, 4C, S5F), similarly to what previous studies overexpressing ZGA regulators in ESCs have shown (Eckersley-Maslin et al. 2019; De Iaco et al. 2019; De Iaco et al. 2017; Hendrickson et al. 2017).

Of the 230 maternal candidates screened, 44 were identified as inducers of a ZGA-like response in ESCs (Fig. 3A-C, 3E). Importantly, amongst these positive hits, we found factors that had been previously described as ZGA regulators, namely Dppa2 (Eckersley-Maslin et al. 2019; De Iaco et al. 2019; Yan et al. 2019), Yap1 (Yu et al. 2016), Gata3 (Eckersley-Maslin et al. 2019) and Ctcf (Wan et al. 2008), further validating our screen approach. Several of our screen hits, such as Cbx1, Smarca5, Phf1, Gatad2b, Smarca2, Hat1, Gata3 or Npm2, have also been shown to significantly reduce the 2-cell like subpopulation in mouse ESCs after siRNA knock-down (Rodriguez-Terrones et al. 2018), consistent with their activator role of ZGA. Other hits have been implicated in, but not specifically shown to regulate ZGA transcription. For example, the binding motif of Arnt is enriched in distal open chromatin regions of embryos undergoing ZGA (Guo et al. 2017).

Consistent with our findings that Smarca5 induces ZGA-like transcription, Smarca5 localizes to sites of active transcription in zygotes and its zygotic RNAi-mediated knock-down led to some defects in ZGA (Torres-Padilla & Zernicka-Goetz 2006). Moreover, Smarca5 homozygous KO embryos derived from heterozygous crosses arrest during preimplantation development (Stopka et al. 2011). DNase-seq analysis of early embryos have shown that transcriptionally inactive genes can have accessible promoters before they become active at later developmental stages (F. Lu et al. 2016). We hypothesize that Smarca5 facilitates ZGA by maintaining an open chromatin state of ZGA gene promoters, and it is not until Dppa2 localizes to the nucleus in the two-cell stage that ZGA is triggered (Fig. 5B-G). Excitingly, the majority of our screen hits, including Patz1 which we independently validated, have not been previously linked to ZGA, thus demonstrating the power of our approach for identifying novel regulators of ZGA. This included Smarca2 and Arid1a (Fig. 3A, 3E), members of the SWI/SNF chromatin remodelling complex. Notably, Smarca5 KO led to a reduction, but not complete elimination, of ZGA-like gene expression in ESCs (Fig. 5E), suggesting there may be functional redundancy amongst chromatin remodellers in the early embryo at this crucial time of development.

In summary, we conclude that our CRISPRa single-cell transcriptomic screen has unravelled novel positive regulators of ZGA. Our data and the candidates identified open up many exciting new avenues for *in vivo* experiments testing the functional requirements, interdependencies and redundancies between these new regulators of ZGA-like expression. Furthermore, our CRISPRa followed by scRNA-seq screening method can be broadly applied in other biological contexts to systematically understand transcriptional regulation at the cellular level and identify positive regulators of key transcriptional programmes in a large-scale and high-throughput manner.

## Materials and Methods

### Cloning

All sgRNAs used in this study were previously designed (Joung et al. 2016) to target the 180 bp region upstream of the target gene TSS. Sequences are provided in Table S1.

The CROP-sgRNA-MS2 lentiviral backbone was synthesized by VectorBuilder by adapting the CROP-seq vector (Datlinger et al. 2017) with the following modifications: 1) the sgRNA scaffold sequence contains two MS2 loops that allow recruitment of MS2-p65-HSF1 in SAM ESCs; and 2) a fluorescent mCherry marker was included downstream of the Eif1a promoter and linked through T2A to a puromycin resistance cassette, allowing assessment of the multiplicity-of-infection (MOI) by FACS and antibiotic selection of the cells (Fig. S1I).

For individual sgRNA cloning in pilot and validation experiments, two oligos were synthesized per sgRNA (Sigma Aldrich), one containing the target sequence (Table S1) with a “CACCG” flank at the 5’ end and the other one synthesized as the reverse complementary sequence to the target sequence and flanked by “AAAC” at the 5’ end and by a “C” at the 3’ end. Each oligo pair was annealed using T4 Polynucleotide Kinase (PNK) enzyme (NEB, M0201S) and then cloned into the sgRNA(MS2)_puro backbone (Addgene 73795) or into the in-house built CROP-sgRNA-MS2 backbone (Fig. S1I) by a Golden Gate reaction using BsmBI enzyme (Thermo Fisher Scientific, ER0451) and T7 ligase (NEB, M0318S). The product from the Golden Gate reaction was transformed into Stbl3 competent cells (Thermo Fisher Scientific, C737303). Between 2 to 3 colonies were picked per sgRNA and verified by Sanger sequencing.

For cloning the 475 sgRNA library into the CROP-sgRNA-MS2 backbone, first, oligos containing the sgRNA target sequence, a 5’ end 26 base-pair (bp) flanking region complementary to the U6 promoter (TATCTTGTGGAAAGGACGAAACACCG) and a 3’ end 35bp flanking region complementary to the sgRNA scaffold sequence (GTTTAAGAGCTAGGCCAACATGAGGATCACCCATG) were synthesized by Twist Bioscience. This oligo library was then amplified and cloned using Gibson assembly, as previously described (Joung et al. 2016), by VectorBuilder. Library coverage was estimated to be >11,000 folds by colony count of diluted transformations. 150 bp paired-end sequencing was performed on Illumina HiSeq4000 to analyse sgRNA representation in the library (Fig. S1J).

cDNA-GFP constructs were cloned by gateway cloning as previously described (Eckersley-Maslin et al. 2019).

### Cell culture, lentiviral transduction, cDNA transient transfections, flow-cytometry and sorting

SAM mouse embryonic stem cells (ESCs) were generated by lentiviral transduction of lenti dCas9-VP64_Blast (Addgene 61425) and lenti MS2-p65-HSF1_Hygro (Addgene 61426) into E14 mouse ESCs, followed by antibiotic selection and manual subcloning. Clones were checked by genomic PCR and one clone was used for consecutive experiments. Dppa2 knock-out (KO) cells were previously described in (Eckersley-Maslin et al. 2019) and Smarca5 (also known as Snf2h) KO cells in (Barisic et al. 2019). All mouse embryonic stem cells (ESCs) were grown under serum/LIF conditions: DMEM (Gibco, 11995-040), 15% fetal bovine serum, 1 U/ml penicillin - 1 mg/ml streptomycin (Gibco, 15140-122), 0.1 mM nonessential amino acids (Gibco, 11140-050), 4 mM GlutaMAX (Gibco, 35050-061), 50 μM β-mercaptoethanol (Gibco, 31350-010), and 103 U/ml LIF (Stem Cell Institute, Cambridge), and cultured at 37 °C in 5% CO_2_ on gelatinized tissue-culture plates. The media was refreshed every day and the cells passaged every other day with Trypsin EDTA (Thermo Fisher Scientific, 25200056). HEK293T cells were grown in D10 media: DMEM (Gibco, 11995-040), 10% fetal bovine serum, 1 U/ml penicillin - 1 mg/ml streptomycin (Gibco, 15140-122) and cultured at 37 in 5% CO_2_ on T175 tissue culture flasks or 100mm tissue culture plates. The media was refreshed every other day and the cells passaged every three days with Trypsin EDTA (Thermo Fisher Scientific, 25200056).

For lentiviral particle production, 3.5 million HEK293T cells were first seeded into 100mm tissue culture plates 24h before transfection. Next, they were cotransfected with 3.5 μg of pMD2.G (Addgene 12259), 6.5 μg of psPAX2 (Addgene 12260) and 10μg of the lentiviral vector of interest: dCas9-VP64_Blast (Addgene 61425), MS2-p65-HSF1_Hygro (Addgene 61426), sgRNA(MS2)_puro (Addgene 73795) cloned with an individual sgRNA, or CROP-sgRNA-MS2 cloned with an individual sgRNA. A single-tube reaction mix was prepared for each transfection, containing the three lentiviral plasmids and 60 μl of TransIT Reagent (Mirus Bio, 2700) diluted in 1.5 ml of Opti-MEM (Gibco, 31985), which was subsequently added drop-wise into the cells containing 8.5 mL of fresh media, following manufacturer’s instructions. 48h later, 10 mL of virus supernatant were harvested by filtering through a 0.45 um filter (Sartorius, 16533) and supplemented with 8μg/ml polybrene (Millpore, TR-1003-G). For sgRNA-expressing lentivirus from the CROP-sgRNA-MS2 backbone, the 10 mL of viral supernatant were concentrated with LentiX Concentrator (Takara, 631231) due to the lower viral titer (Datlinger et al. 2017).

The lentiviral 475 sgRNA library was packaged by VectorBuilder: the plasmid library was co-transfected with a proprietary envelop plasmid encoding VSV-G and packaging plasmids encoding Gag/Pol and Rev into packaging cells. After a short incubation period, the supernatant was collected, followed by the removal of cell debris by centrifugation, filtration and PEG concentration of the viral particles.

To measure lentiviral titer, HEK293T cells were transduced with lentivirus diluted from the stock and then, a quantitative reverse transcription PCR-based approach was used to quantify the average number of integration events of the proviral genome per host genome.

Individual sgRNAs were transduced into SAM ESCs by direct supplementation of the lentivirus into the medium for 24 hours. 48 hours after transduction, 1 μg/ml puromycin was added to the medium for selection of transduced cells. Cells were selected and passaged for 8 days after addition of puromycin before harvesting for bulk or 10X single-cell RNA-sequencing library preparation.

The lentiviral library of 475 sgRNAs cloned into the CROP-sgRNA-MS2 backbone was transduced in triplicate into SAM ESCs by direct supplementation of the concentrated lentivirus into the medium for 24 hours. The transductions were done at <0.1 multiplicity-of-infection (MOI) (Fig. S2A) into 5 million SAM ESCs, achieving a representation >1000 cells / sgRNA, considering that <10% of the cells were transduced. Two days after transduction, mCherry expression was analysed by flow-cytometry on BD LSR Fortessa and selection with 1 μg/ml puromycin started afterwards. Cells were selected and passaged for 8 days after addition of puromycin before harvesting for 10X single-cell RNA-sequencing library preparation.

cDNA-eGFP constructs were transfected into pre-plated E14, Dppa2 KO, Smarca5 KO or wild-type counterparts mouse ESCs in triplicates using Lipofectamine 2000 Reagent Opti-MEM (GIBCO, 31985) (Thermo Fisher Scientific, 11668019), following manufacturer’s instructions. Cells were grown for 48 hours before GFP^+^ FACS sorting on BD Influx High-Speed Cell Sorter or analysis on BD LSR Fortessa.

### Preparation of 10X 3’ single-cell RNA-sequencing libraries

A single cell suspension was loaded into the 10X Chromium device and libraries were prepared using the 10X V2 Single Cell 3′ Solution kit (10X Genomics) following manufacturer’s instructions. In the pilot test, the following samples were loaded each in a lane of the 10X chip Chromium Controller: E14 ESCs, SAM ESCs transduced with the non-targeting control 461, SAM ESCs transduced with the

MERVL LTR sgRNAs 459 and 460 individually and pooled at the time of sequencing, and SAM ESCs transduced with the Zscan4 sgRNAs 457 and 458 individually and pooled at the time of sequencing (see Table S1 for sgRNA sequences). Each 10X library was sequenced on an Illumina HiSeq4000 lane with 75 cycles for read 1, 75 cycles for read 2 and 8 cycles i7 sample index. 1956, 2045, 2233, and 2362 cells were captured, respectively, for each sample based on number of cell barcodes detected, before quality control, with a total of 27,998 genes detected.

In the single-cell dataset of SAM ESCs transduced with the 475 sgRNA library, each transduction replicate was loaded across a full 10X chip Chromium Controller (8 lanes), with 20,000 cells per lane. Each 10X library was sequenced on an Illumina HiSeq4000 lane with 26 cycles for read 1, 98 cycles for read 2 and 8 cycles i7 sample index. A total of 114,866 cells were captured for replicate 1, 118,646 cells for replicate 2 and 107,591cells for replicate 3 based on number of cell barcodes detected, before quality control, which, after merging transduction replicates, resulted in a dataset of 341,103 cells with a total of 27,998 genes detected before quality control processing.

Amplicon sgRNA PCRs were performed for each of the 24 full length 10X cDNA samples of SAM ESCs transduced with the 475 sgRNA library, as previously described (Hill et al. 2018). Briefly, 10 ng of full length 10X cDNA were used as starting material and each round of PCR amplification with the primers described in (Hill et al. 2018) was monitored by KAPA SYBR (KR0389) to avoid overcycling. After multiplexing, these enrichment libraries were sequenced across two lanes of the Illumina Hiseq2500 Rapid Run, with 27 cycles for read 1, 267 cycles for read 2 and 8 cycles for i7 sample index.

### Processing and analysis of 10X 3’ single-cell RNA-sequencing data from pilot test

*Processing, quality control and gene expression quantification:* All 10X scRNA-seq data was processed with the default CellRanger v2.1 pipeline (Zheng et al. 2017) for mapping to the mm10 mouse genome assembly. Gene counts were further analysed with scanpy (Wolf et al. 2018). For quality control of the samples from the pilot test after individual transductions with MERVL LTRs and Zscan4 sgRNAs, cells with less than 15,000 UMI counts and/or less than 4,000 detected genes, cells with more than 40,000 UMI counts and/or more than 6,500 detected genes, and cells with more than 5% of UMI reads coming from mitochondrial genes were discarded (Fig. S1B-D). After this quality control, 1,138 E14 ESCs, 687 SAM ESCs transduced with the non-targeting control 461, 1227 SAM ESCs transduced with the MERVL LTR sgRNAs 459 and 460 and 899 SAM ESCs transduced with the Zscan4 sgRNAs 457 and 458 were retained for analysis. A gene was considered for downstream analysis if it was detected (UMI count >0) in at least 10 cells that passed the quality control filter across the full dataset. The final dataset consisted of 16,498 genes across 3,951 cells. The number of UMIs for each cell and gene were adjusted by the library size in each cell, dividing by the total number of UMIs per cell. Gene expression levels were obtained as log-transformed adjusted UMI counts, scaled by a factor of 10,000. For quantification of MERVL repeat elements, see “repeat element quantification” section below.

*Power estimation (number of cells to be sequenced per sgRNA):* From the pilot experiment where SAM ESCs were transduced with sgRNAs targeting MERVL LTRs and Zscan4 cluster promoters (Fig. S1E-F), a power estimation was made to determine the number of single cells required to be sequenced per sgRNA in the CRISPRa screen to detect a ZGA-like signature in a positive hit. We used a qualitative two-tailed Fisher’s exact test considering that the percentage of cells that expressed ZGA-like transcripts was 2.04% in cells transduced with a non-targeting sgRNA control and 8.56% in cells transduced with a positive inductor (e.g. MERVL LTR sgRNAs) (Fig. S1G). The test returned a sample size of 399 cells per sgRNA to detect a ZGA-like signature in a positive hit with an adjusted p-value < 0.00032 and 0.8 power.

### Processing and analysis of 10X 3’ single-cell RNA-sequencing data from the primary CRISPRa screen

*Processing, quality control and gene expression quantification:* All 10X scRNA-seq data was processed with the default CellRanger v2.1 pipeline (Zheng et al. 2017) for mapping to the mm10 mouse genome assembly. Gene counts were further analysed with scanpy (Wolf et al. 2018). To keep only high-quality cells, we filtered out cells with less than 4,000 UMI counts and/or less than 1,600 detected genes, cells with more than 20,000 UMI counts and/or more than 5,000 genes, and cells with more than 5% of UMI reads coming from mitochondrial genes (Fig. S2B-D). After filtering, 109,061 cells were retained from replicate 1, 118,646 cells from replicate 2 and 107,591 cells from replicate 3. Next, we assigned a sgRNA to each cell using the amplicon sgRNA libraries (see Assignment of sgRNAs to cells below) and we discarded cells that were not uniquely assigned to one sgRNA, resulting in 71,047 cells in replicate 1, 71,188 cells in replicate 2 and 61,729 cells in replicate 3, which corresponds to 203,894 cells in total across all replicate sets (Fig. 1B). A gene was considered for downstream analysis if it was detected (UMI count >0) in at least 10 out of the 203,894 cells that passed filtering. The final dataset after quality control consisted of 20,690 genes. The number of UMIs for each cell and gene were adjusted by the library size in each cell, dividing by the total number of UMIs per cell. Gene expression levels were obtained as log-transformed adjusted UMI counts, scaled by a factor of 10,000. For PCA analysis (Fig. 1C, S2I-J), 965 highly variable genes were selected, as implemented in scanpy (Wolf et al. 2018) with minimum mean of 0.01, max mean of 5 and minimum dispersion of 0.5.

*Assignment of sgRNAs to cells:* Using the amplicon sgRNA libraries, the potential sgRNA sequence (nucleotides 24-43 of the read) was compared to the collection of designed sgRNAs. By taking only exact matches to the white list of sgRNAs, the majority of 475 sgRNA sequences were recovered (470-474, variable from one library to another) and 16% of reads on average (15.3%-16.8% for different libraries) were left unassigned to a sgRNA. To correct for sequencing errors, we allowed a minimum edit distance (Levenshtein distance – 4 edits) between any two sequences of the designed sgRNAs as well as the CROP-sgRNA-MS2 vector sequence surrounding the potential sgRNA in the read. For the reads left unassigned to a sgRNA at this stage, if there was a sgRNA sequence within Levenshtein distance of 1 or 2 and if the upstream 23 nucleotides and downstream 23 nucleotides matched the CROP-sgRNA-MS2 vector sequence with up to 4 edits each, the respective sgRNA was assigned to the read. After this correction procedure, approximately 2% of reads were left unassigned. Cell barcodes detected in the amplicon libraries were then matched with barcodes detected in the regular 10X scRNA-seq libraries. Out of the 317,847 cells that passed quality control across the three transduction replicates in the regular 10X libraries, 249,767 cell barcodes were captured in the amplicon sgRNA libraries (85,993 in replicate 1, 86,671 in replicate 2 and 77,103 in replicate 3). A sgRNA was assigned to a cell if more than 90% of all the amplicon reads containing the sgRNA had the same cell barcode, with standard error of binomial proportion of less than 10% (e.g. more than 8 reads if all the barcodes are associated with the same sgRNA, 13 reads of the same sgRNA if there were more than one sgRNA for a cell barcode, etc). The following table illustrates cell numbers and percentages for each assignment in each replicate (see also Fig. 1B):

| Replicate | Total number of cells | Assignment | Number of cells | Percentage |
| --- | --- | --- | --- | --- |
| 1 | 85,933 | No sgRNA | 397 | 0.46 |
|  | 85,933 | Unique sgRNA | 71,047 | 82.62 |
|  | 85,933 | Two sgRNAs | 3,028 | 3.52 |
|  | 85,933 | Multiple sgRNAs | 11,521 | 13.40 |
| 2 | 86,671 | No sgRNA | 400 | 0.46 |
|  | 86,671 | Unique sgRNA | 71,118 | 82.14 |
|  | 86,671 | Two sgRNAs | 3,210 | 3.70 |
|  | 86,671 | Multiple sgRNAs | 11,873 | 13.70 |
| 3 | 77,103 | No sgRNA | 381 | 0.49 |
|  | 77,103 | Unique sgRNA | 61,729 | 80.06 |
|  | 77,103 | Two sgRNAs | 3,084 | 4.00 |
|  | 77,103 | Multiple sgRNAs | 11,909 | 15.45 |

*Repeat elements quantification:* All occurrences in the genome of repeat sequences from 12 repeat families (LINE-1, LINE-2, ERV1, ERVK, MERVL, Major satellites, Minor satellites, ribosomal RNA, SINE Alu B1, SINE B2, SINE B4, Telomeric repeats), with each respective genomic locations were downloaded from the UCSC table browser (RepeatMasker, mm10, Nov 2018), concatenated and treated as a reference genome to map the reads discarded by CellRanger pipeline, due to mapping to multiple regions, using SAMtools (H. Li et al. 2009) and BWA (version 0.7.17-r1188, default parameters) (H. Li & Durbin 2009). The following number of reads were discarded by CellRanger pipeline in each transduction replicate: 253,330,874 reads in replicate 1, 276,401,843 reads in replicate 2, 242,863,617 reads in replicate 3, out of which 38,792,331 (15.31%) in replicate 1, 37,285,351 (13.49%) in replicate 2 and 25,837,444 (10.64%) mapped to repeat elements. LINE-2 elements and Minor satellite repeats were discarded from downstream analysis due to inefficient mapping (see table below). Reads sharing a unique molecular identifier and a cell barcode were then collapsed in order to get an estimate of the number of molecules for each repeat family in every cell (Fig. S3A-B).

| Replicate | Repeat family | Number of mapped reads |
| --- | --- | --- |
| 1 | LINE-1 | 15,930,866 |
|  | LINE-2 | 56 |
|  | ERV1 | 1,049,021 |
|  | ERVK | 14,783,904 |
|  | MERVL | 1,275,515 |
|  | Major Satellites | 305,524 |
|  | Minor Satellites | 3 |
|  | Ribosomal RNA (rRNA) | 773 |
|  | SINE Alu B1 | 398,331 |
|  | SINE B2 | 4,583,807 |
|  | SINE B4 | 464,024 |
|  | Telomeric repeats | 447 |
| 2 | LINE-1 | 14,790,530 |
|  | LINE-2 | 86 |
|  | ERV1 | 1,159,969 |
|  | ERVK | 13,299,950 |
|  | MERVL | 1,382,105 |
|  | Major Satellites | 318,816 |
|  | Minor Satellites | 2 |
|  | Ribosomal RNA (rRNA) | 846 |
|  | SINE Alu B1 | 457,910 |
|  | SINE B2 | 5,341,151 |
|  | SINE B4 | 533,479 |
|  | Telomeric repeats | 507 |
| 3 | LINE-1 | 9,769,600 |
|  | LINE-2 | 43 |
|  | ERV1 | 952,484 |
|  | ERVK | 8,865,749 |
|  | MERVL | 1,049,799 |
|  | Major Satellites | 173,969 |
|  | Minor Satellites | 4 |
|  | Ribosomal RNA (rRNA) | 551 |
|  | SINE Alu B1 | 370,782 |
|  | SINE B2 | 4,226,418 |
|  | SINE B4 | 427,651 |
|  | Telomeric repeats | 394 |

*Multi-omics factor analysis (MOFA) model:* A hierarchical Bayesian model, as implemented in an extension of MOFA (unpublished), was trained on two views: first, the set of 965 highly variable coding genes, and second, the expression levels of eight repeat families (LINE-1, ERV1, ERVK, MERVL, Major satellites, SINE Alu B1, SINE B2 and SINE B4). The ribosomal RNA (rRNA) and telomeric repeats were excluded from the model due to low detection rate (Fig. S3A-B). The cell-to-sgRNA assignment to group cells based on sgRNA expression was provided to the model in order to take advantage of group-wise sparsity of the model (Fig. 2A). Upon interpreting the first five factors using their top loadings of variance explained (Table S4, Fig. 2B, 2D, S3C, S3G), factor 3 was interpreted as a ZGA-like factor due to the top loadings (both coding genes and MERVL repeat) being genes expressed at the time of ZGA (Fig. 2B-D).

*Identification of screen hits:* MOFA factor 3 was used to reconstruct the expression dataset for each group of cells to calculate the variance explained by factor 3 per group. Average fraction of expression variance for coding genes and repeat elements was then used to rank the 460 targeting sgRNAs (Table S1), with higher percent of variance explained being attributed to sgRNAs with more potency to induce a ZGA-like transcriptional signature. For hit calling, the average and one standard deviation of the fraction of expression variance explained by MOFA factor 3 for cells expressing the fifteen different non-targeting sgRNA controls was estimated as a background rate, and sgRNAs ranking higher than the standard deviation above the mean were considered hits (Fig. 3A, Table S1). Higher fraction of variance explained by MOFA factor 3 correlates with higher factor 3 values (Fig. S4A) and hence higher expression of ZGA-like genes (Fig. 3B, 3C).

*Differential gene expression analysis:* For every targeting sgRNA in the screen, respective cells were compared to the set of cells with non-targeting sgRNA controls. A generalised linear model (glm) as implemented in EdgeR was fitted for every gene, and a likelihood ratio test was used to estimate the effect of the targeting sgRNA on the gene’s level of expression (Robinson et al. 2010).

### Preparation of bulk RNA-sequencing libraries

RNA was isolated using RNeasy Mini kit (Qiagen, 74104) and treated with DNaseI (Thermo Fisher Scientific, EN0521) following manufacturer’s instructions.

For SAM and E14 ESCs, opposite strand-specific total RNA libraries (ribozero) were made using 1 µg DNase-treated RNA using the Sanger Institute Illumina bespoke pipeline and sequenced at 100 bp paired-end on the Illumina HiSeq 2500 Rapid Run platform. For CRISPRa (see Table S1 for sgRNAs used) and cDNA validation samples, opposite strand-specific polyA RNA libraries were made using 1 µg DNase-treated RNA using the Sanger Institute Illumina bespoke pipeline and sequenced at 50 bp single-end on Illumina HiSeq4000.

All bulk RNA-sequencing experiments were performed in three independent replicates, except SAM22 mESCs for which two replicates were prepared.

### Processing and analysis of bulk RNA-sequencing datasets

For processing of all RNA-sequencing data, including those generated in this study but also re-analysis of (Xue et al. 2013; Deng et al. 2014; Barisic et al. 2019), raw FastQ data were trimmed with Trim Galore (v0.6.1, default parameters) and mapped to the mouse mm10 genome assembly using Hisat2 v2.0.5, as guided by known splice sites taken from Ensembl v68. Hits were filtered to remove mappings with MAPQ scores <20. Data were quantitated at mRNA level using the RNA-seq quantitation pipeline in SeqMonk software (www.bioinformatics.babraham.ac.uk/projects/seqmonk/) with strand specific quantification using mRNA probes. For alignments to dCas9-VP64 and MS2-p65-HSF1 exogenous integrations, we constructed an artificial genome and integrated to mm10 to quantify their expression in relation to the whole transcriptome (Fig. S1A).

For alignments to repetitive regions in the genome we constructed artificial repeat genomes. Repeat annotations were downloaded from the UCSC table browser (RepeatMasker, mm10, Nov 2018). Sequences of the list of repeat element instances were stitched together separated by ‘NNNNN’ to create repeat specific genomes. Trimmed reads from each sample were aligned against all individual repeat genomes using Bowtie2 (v2.3.2). Values given are cumulative reads mapping to a specific repeat group as percentage of the total read count.

Differentially-expressed genes were determined using EdgeR (FDR<0.5). Individual CRISPRa of Dppa2, Smarca5 and Patz1 replicate sets were compared to non-targeting controls 461 and 462 for differential gene expression. Differential gene expression for Dppa2-GFP^+^, Smarca5-GFP^+^ and Patz1-GFP^+^ was done against GFP^+^ only controls. The genes that were differentially expressed both by CRISPRa and cDNA overexpression for each target gene were used in Fig. 4D. Whole transcriptome correlations were calculated using Pearson correlation coefficient on replicate sets.

### Mice and embryo collection

All mice used in this study were C57BL/6 and were bred and maintained in the Babraham Institute Biological Support Unit. All procedures were covered by a project license (to WR) under the Animal (Scientific Procedures) Act 1986, and are locally regulated by the Babraham Institute Animal Welfare, Experimentation, and Ethics Committee. Embryos were collected from C57BL/6 females after superovulation and mating to C57BL/6 males. Zygotes were collected on the day of plugging and two-cell embryos one day after plugging.

### Immunofluorescence, imaging and co-localization quantification

Embryos were collected in M2 medium (Sigma-Aldrich, MR-015P-5F) containing hyaluronidase (Sigma, H2126), and washed in M2 droplets to remove the cumulus cells. After fixation with 4% PFA (Polysciences, Inc., 18814) for 10 minutes at room temperature, embryos were permeabilised with 0.5% TritonX-100 in PBS for 1 hour and blocked with 1% BSA in 0.05% Tween20 in PBS (BS) for 1hour. Primary antibodies were diluted in BS and incubated for 1 hour, followed by 1hour wash in BS. Next, secondary antibodies diluted in BS were incubated for 45 minutes, followed by 30 minutes to 1hour wash in 0.05% Tween20 in PBS. All incubations were performed at room temperature. Primary antibodies and dilutions used were: Smarca5 (ab72499) 1:100 and Dppa2 (mab4356) 1:200. All secondary antibodies were Alexa Fluor conjugated (Molecular Probes - anti-rabbit AF 568 and anti-mouse AF 488) and diluted 1:1000. DNA was counterstained with 5µg/mL DAPI in PBS. Embryos were mounted in fibrin clots. Single optical sections were captured with a Zeiss LSM780 microscope (63x oil-immersion objective) and the images pseudo-colored using ImageJ2. For visualization, images were corrected for brightness and contrast, within the recommendations for scientific data. Fluorescence co-localization analysis was performed with Volocity 6.3 (Improvision) in 10 zygotes and 10 two-cell embryos. Pearson’s correlation coefficient between Smarca5 and Dppa2 signals were calculated in the area corresponding to the pronuclei or nuclei. The pronuclei in zygotes and nuclei of each blastomere in two-cell embryos were measured separately, with values comparable within the same embryo.

### Quantitative reverse transcription PCR (qPCR)

RNA was isolated using RNeasy Mini kit (Qiagen, 74104) and treated with DNaseI (Thermo Fisher Scientific, EN0521) following manufacturer’s instructions. cDNA was synthesized from 0.5-2 μg of DNAaseI-treated RNA using RevertAid First-Strand cDNA Synthesis Kit (Thermo Fisher Scientific, K1622), and diluted 1:10 prior to qPCR. qPCR was performed in technical duplicates using Brilliant III SYBR master mix (Agilent Technologies, 600882) and a CFX384 Touch Real-Time PCR Detection System machine (BioRad). Relative levels of transcript expression were quantified by the comparative CT method with normalisation to Gapdh levels. Primer sequences are available upon request.

### Data representation

Graphs and illustrations were performed with RStudio, SeqMonk, GraphPad Prism, Microsoft Excel and Illustrator software.

## Supplemental Tables Information

**Table S1**: Related to pooled sgRNA library, CRISPRa scRNA-seq dataset with three replicates merged, screen results, and information on sgRNAs used for pilot and validation experiments. The table contains sequence information on the 475 sgRNAs used in the study, target gene, target transcript and sequencing results of the pooled sgRNA plasmid library. It also contains information on the CRISPRa scRNA-seq screen dataset, such as number of cells expressing each sgRNA, expression of the target gene and differential gene expression analyses. Screen results (fraction of expression variance explained by MOFA factor 3, hit rank and whether the sgRNA was considered a hit) are also shown in this table.

**Table S2**: Gene names of ZGA signature. The table contains the gene names of previously identified ZGA genes in (Eckersley-Maslin et al. 2016; Hendrickson et al. 2017; Y. Li et al. 2018). The list is a combination of Table S1 from Eckersley-Maslin et al. 2016, Table S8 from Hendrickson et al. 2017 and Table S1 from Y. Li et al. 2018.

**Table S3**: Related to principal component analysis (PC1). The table contains loading values of 965 highly-variable genes in the CRISPRa scRNA-seq screen dataset for the first two principal components (PC1 and PC2) in tab 1, gene ontology enrichment results of the top 50 loading genes for PC1 in tab 2 and gene ontology enrichment results of the top 50 loading genes for PC2 in tab 3.

**Table S4**: Related to MOFA. The table contains loading values of 965 highly-variable genes in the CRISPRa scRNA-seq screen dataset for MOFA factors 1-5.

## Acknowledgements

The authors thank all members of the Reik and Stegle laboratories for helpful discussions. We also thank Mario Iurlaro for helpful discussions on Smarca5, Paul Datlinger for advice on CROP-seq lentiviral preparation, Felix Krueger and the bioinformatics facility from the Babraham Institute for processing sequencing data and assistance with repeat mapping, Lia Chappell for training in 10X Genomics library preparation, Anne Segonds-Pinchon for statistical advice, the sequencing facilities at Babraham Institute, Sanger Institute and CRUK in Cambridge for high-throughput library preparation and sequencing, and the flow cytometry facility at the Babraham Institute for cell sorting. Smarca5 (Snf2h) KO ESCs were a kind gift from Dirk Schübeler; lenti sgRNA(MS2)_puro backbone (Addgene 73795), lenti dCAS-VP64_Blast (Addgene 61425) and lenti MS2-P65-HSF1_Hygro (Addgene 61426) were a gift from Feng Zhang; pMD2.G (Addgene 12259) and psPAX2 (Addgene 12260) were a gift from Didier Trono. C.A.-C. is supported by a postgraduate award by the Biotechnology and Biological Sciences Research Council (BBSRC). D.B is supported by a Darwin Trust fellowship. I.H.-H. is supported by a Marie Sklodowska-Curie Individual Fellowship (751439). Research in the Reik lab is supported by BBSRC (BB/K010867/1), Wellcome Trust (095645/Z/11/Z) and EU EpiGeneSys. Research in the Stegle lab is supported by the BMBF, the Volkswagen Foundation and the EU (ERC project DECODE).

## Author contributions

C.A.-C, M.A.E.-M. and W.R. conceived and designed the study. C.A.-C. performed and analysed experiments, performed bioinformatic analyses and wrote the original draft. D.B. conceived and performed bioinformatic analyses. I.H.-H analysed the initial pilot test and gave advice and bioinformatic analyses. O.K and F.S performed Dppa2 and Smarca5 immunofluorescence staining. O.S. conceived and supervised bioinformatic analyses. All authors reviewed and edited the manuscript. C.A.-C, M.A.E.-M., O.S. and W.R. supervised the study.

---

**Supplemental Figure 1. Related to Figure 1.**
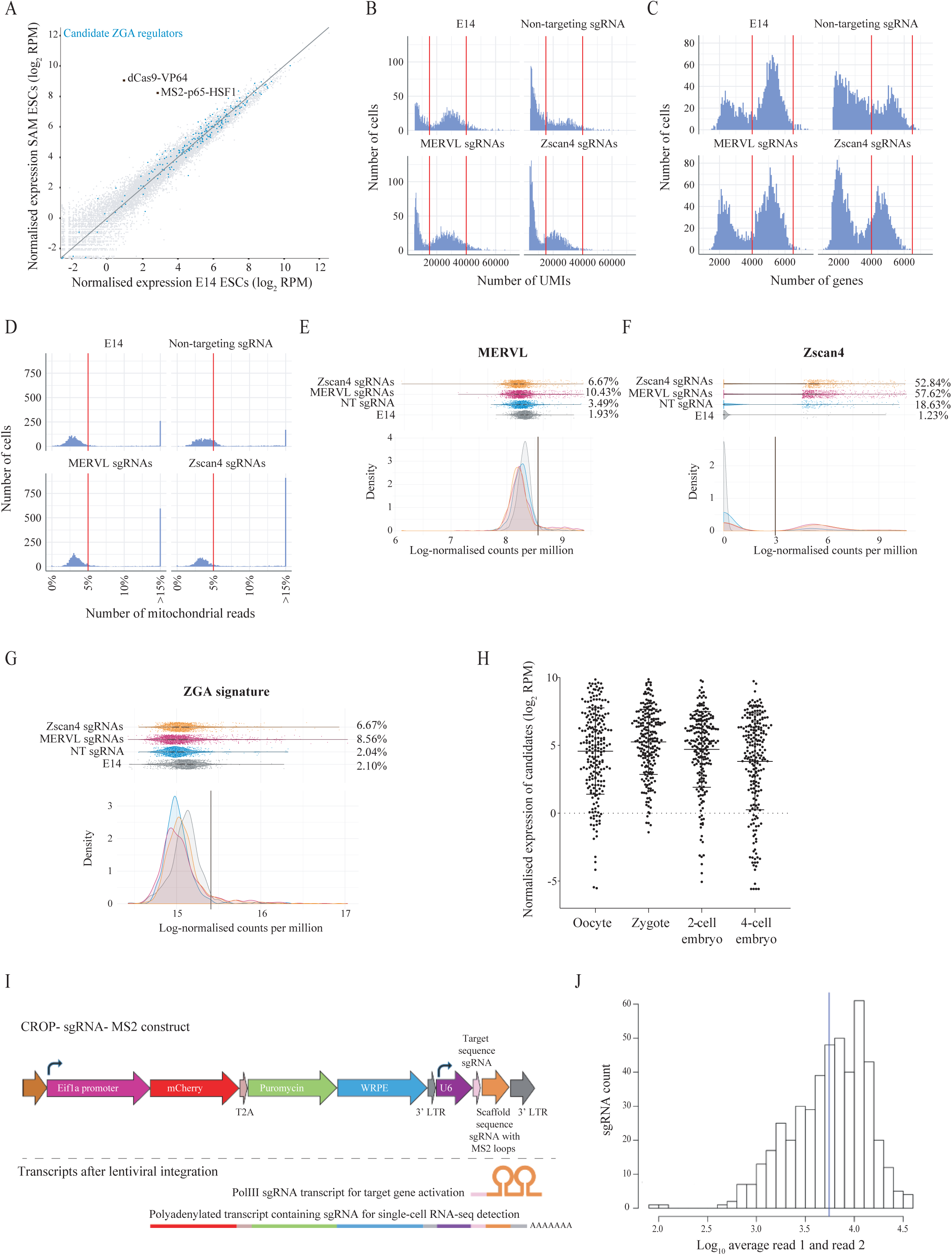
**A)** Scatterplot showing normalised expression levels (log_2_ reads per million; RPM) of SAM ESCs against their parental cell line E14, analysed by RNA-sequencing, highlighting dCas9-VP64 and MS2-p65-HSF1 transcripts in black and screening candidates in blue. **B)** Histograms showing number of unique molecular identifiers (UMIs) for each analysed sample; cells within the red bars (15,000<40,000 UMIs) were retained as quality control (QC)-passing cells. **C)** Histograms showing number of detected genes for each analysed sample; cells within the red bars (4,000<6,500 genes) were retained as QC-passing cells. **D)** Histograms showing the percentage of reads from mitochondrial reads for each analysed sample; cells below the red bars (<5%) were retained as QC-passing cells. **E, F)** Dot-plot (upper panels) and density representation (lower panels) of MERVL **(E)** and Zscan4b/c/d/e/f (sum of all genes) **(F)** log-normalised expression levels in counts per million in untransduced E14 ESCs (grey), SAM ESCs transduced with a non-targeting sgRNA (blue), SAM ESCs transduced with sgRNAs targeting MERVL LTRs (pink) and SAM ESCs transduced sgRNAs targeting Zscan4 gene promoters (orange), analysed by 10X Genomics 3’ single-cell RNA-sequencing. Reported percentages were calculated with the cells above the vertical black line from the total number of cells in each sample. **G)** Dot-plot (upper panels) and density representation (lower panels) of ZGA genes (as described in Table S2) log-normalised expression in counts per million (sum of all genes) in the samples described in **E)** and **F)**. Reported percentages were calculated with the cells above the vertical black line from the total number of cells in each sample. **H)** Dot-plot showing normalised expression levels (log_2_ reads per million; RPM) of the screening candidates in oocytes, zygotes, 2-cell and 4-cell embryos. Data analysed from Xue et al. 2013). **I)** Schematic representation of CROP-sgRNA-MS2 lentiviral vector and transcripts produced upon lentiviral reverse transcription and integration in the host genome. **J)** Histogram showing normalised log_10_ average of read 1 and read 2 from next-generation sequencing analysis of the 475 oligo sgRNA library cloned into the lentiviral backbone described in **I)**; representation of 90% of sgRNAs in the library is within 19.9 folds.

**Supplemental Figure 2. Related to Figure 1.**
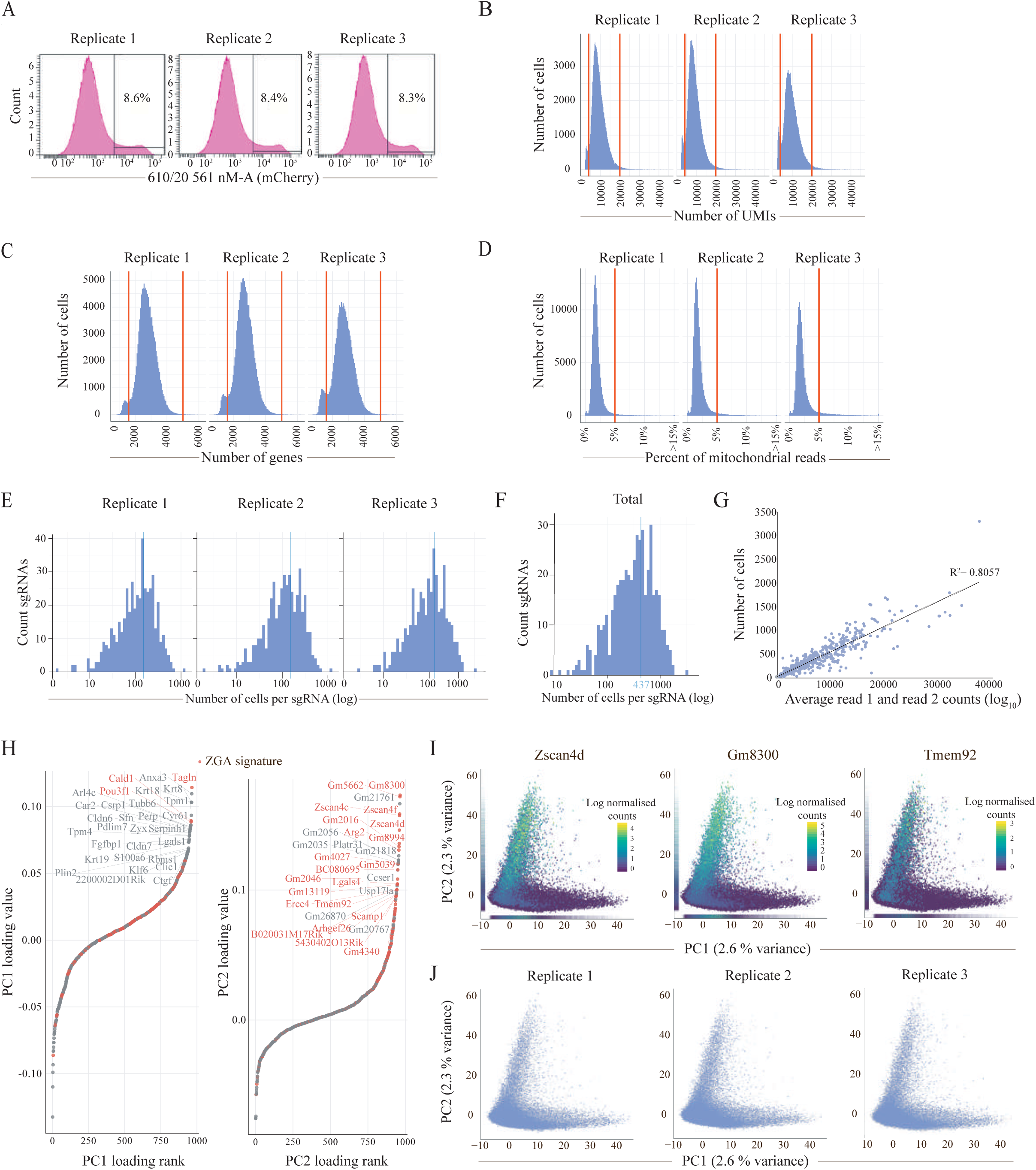
**A)** Histograms showing expression of mCherry analysed by flow cytometry in three replicates of SAM ESCs transduced with the 475 sgRNA lentiviral library at low multiplicity-of-infection (MOI); percentages of cells transduced (expressing mCherry) are shown. **B)** Histograms showing number of unique molecular identifiers (UMIs) for each transduction replicate; cells within the red bars (4,000<20,000 UMIs) were retained as quality control (QC)-passing cells. **C)** Histograms showing number of detected genes for each transduction replicate; cells within the red bars (1,600<5,000 genes) were retained as QC-passing cells. **D)** Histograms showing the percentage of reads from mitochondrial reads for each transduction replicate; cells below the red bars (<5%) were retained as QC-passing cells. **E, F)** Histogram of the number of cells expressing a given unique sgRNA in the 475 sgRNA library for each transduction replicate **(E)** and in the combined dataset across all the three replicates **(F)**. The number of cells per sgRNA is depicted in logarithmic scale. The vertical blue line in **F)** represents the average of 437 cells per sgRNA in the dataset. **G)** Scatterplot between the number of cells expressing a given sgRNA in the scRNA-seq activation screen data (y axis) versus the normalised representation of the sgRNA in the cloned oligo library before transduction (log_10_ average of read 1 and read 2-x axis). The sgRNA representation and the number of cells expressing a given sgRNA are correlated (R^2^ = 0.8057). **H)** Genes ranked by the loadings of the first principal component (PC1; left) and the second principal component (PC2; right), highlighting in red previously know ZGA genes (as described in Table S2, see also Table S3 for gene loading values). **I)** Principal component analysis of all cells that passed quality control, displaying a scatterplot of the first two principal components (PC1 versus PC2) with cells coloured by the expression of the ZGA markers Zscan4d, Gm8300 and Tmem92. Marginal distributions of PC1 and PC2 values are displayed as rug plots along the respective axis. **J)** Reproducibility across the three transduction replicates: principal component analysis of cells in the combined dataset, highlighting in each panel the cells from each transduction replicate that contribute to the analysis.

**Supplemental Figure 3. Related to Figure 2.**
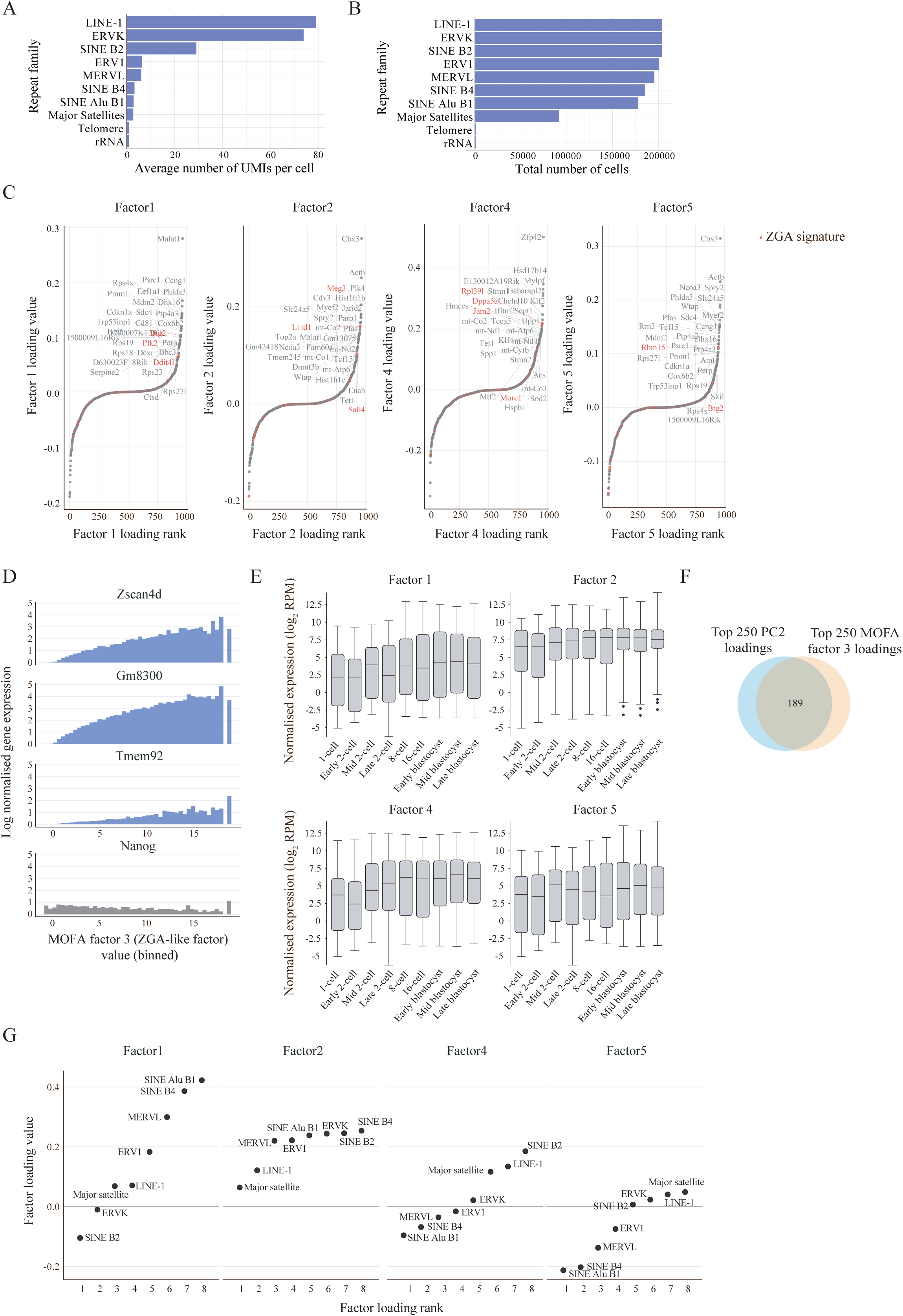
**A)** Average number of UMIs per cell in the combined dataset after quality control for each of ten repeat families analysed. **B)** Total number of cells in the combined dataset after quality control for which each repeat family is detected (UMI>0), out of a total of 203,894 cells analysed. **C)** Coding genes ranked by their loadings of MOFA factors 1, 2, 4 and 5, highlighting in red previously know ZGA genes (as described in Table S2, see also Table S4 for gene loading values). **D)** Normalised expression levels in logarithmic scale for the ZGA markers Zscan4d, Gm8300 and Tmem92 (blue) and the unrelated gene Nanog (grey), for different quantiles of MOFA factor 3 (ZGA-like factor) values, showing ZGA markers contribute to the variance explained in this MOFA factor. **E)** Box-whisker plots showing normalised expression levels (log_2_ reads per million; RPM) for the top 50 gene loadings of MOFA factors 1, 2, 4 and 5 during preimplantation development (data analysed from Deng et al. 2014) (see Table S4 for gene loadings). **F)** Venn diagram showing overlap between the top 250 loadings of component 2 identified using conventional principal component analysis (PC2) and the top 250 loadings of MOFA factor 3 (ZGA-like factor) (see Table S4 for gene loadings). **G)** Repeat element families ranked by their loadings of MOFA factors 1, 2, 4 and 5.

**Supplemental Figure 4. Related to Figure 3.**
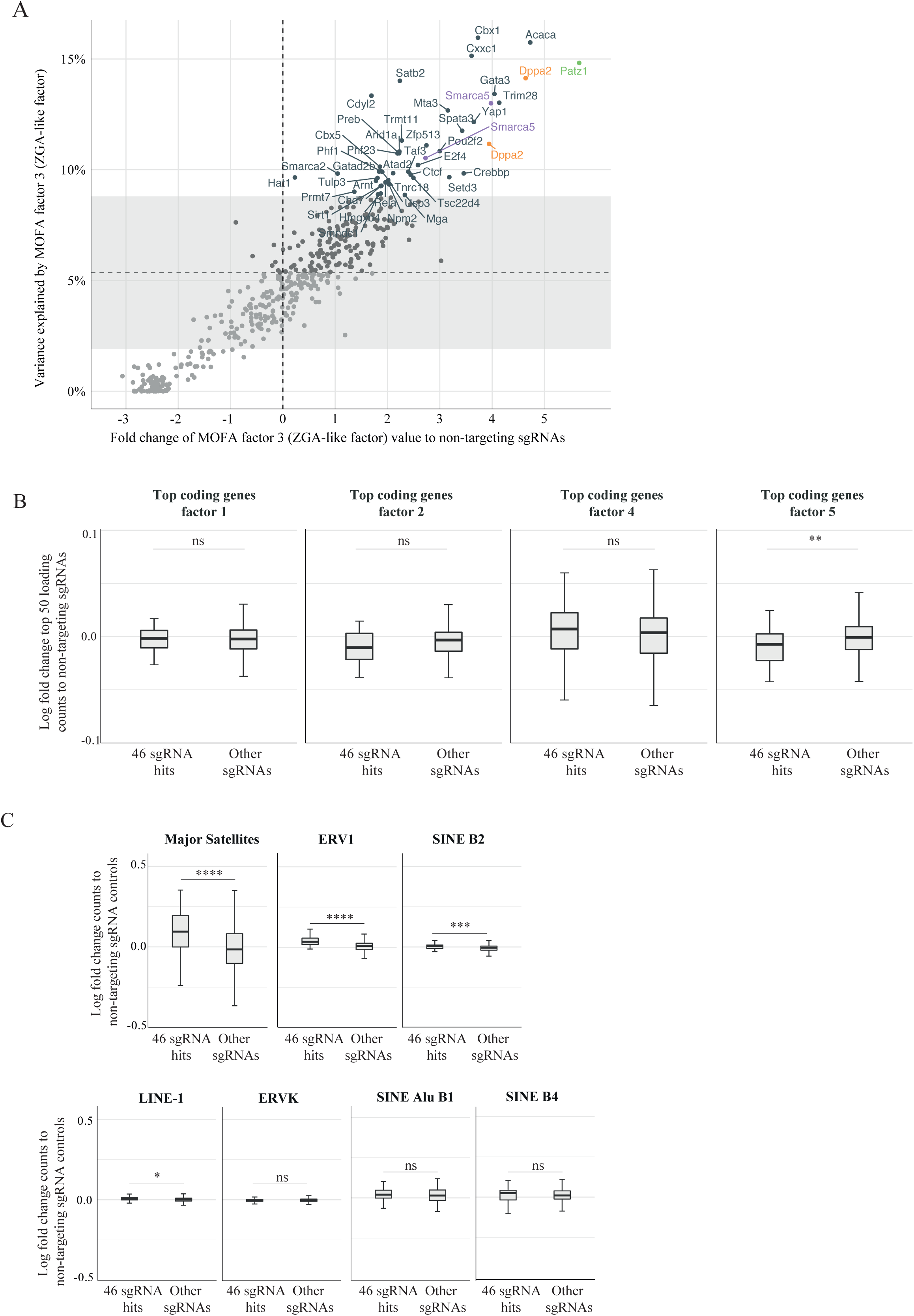
**A)** Scatterplot between the fraction of expression variance (average of coding genes and repeat elements) explained by MOFA factor 3 (ZGA-like factor) for each targeting sgRNA (y axis) and the fold change of the average MOFA factor 3 values for cells expressing each targeting sgRNA to the average MOFA factor 3 values for cells expressing non-targeting sgRNA controls (x axis). The fraction of expression variance explained by the MOFA factor 3 for the fifteen non-targeting sgRNA controls is depicted in the background as mean (horizontal dashed line) and plus and minus one standard deviation (shaded area). The vertical dashed line at *x* = 0 is shown to depict positive fold changes above this value. The names of the target genes for sgRNAs identified as hits (see Table S1) are shown, with Dppa2 (orange), Smarca5 (purple) and Patz1 (green) sgRNA hits highlighted. **B)** Box-whisker plots showing log fold change expression the top 50 loading genes of MOFA factors 1, 2, 4 and 5 in cells expressing the 46 sgRNA hits and cells expressing other targeting sgRNAs, compared to cells expressing non-targeting sgRNA controls. Expression is quantified in normalised counts. (**: p-value <0.01, ns: non-significant; Mann-Whitney two-tailed t-test). **C)** Box-whisker plots showing log fold change of normalised counts of different repeat families counts in cells expressing the 46 sgRNA hits and cells expressing other targeting sgRNAs, compared to cells expressing non-targeting sgRNA controls (****: p-value < 0.0001, ***: p-value <0.001, *: p-value <0.05, ns: non-significant; Mann-Whitney two-tailed t-test).

**Supplemental Figure 5. Related to Figure 4.**
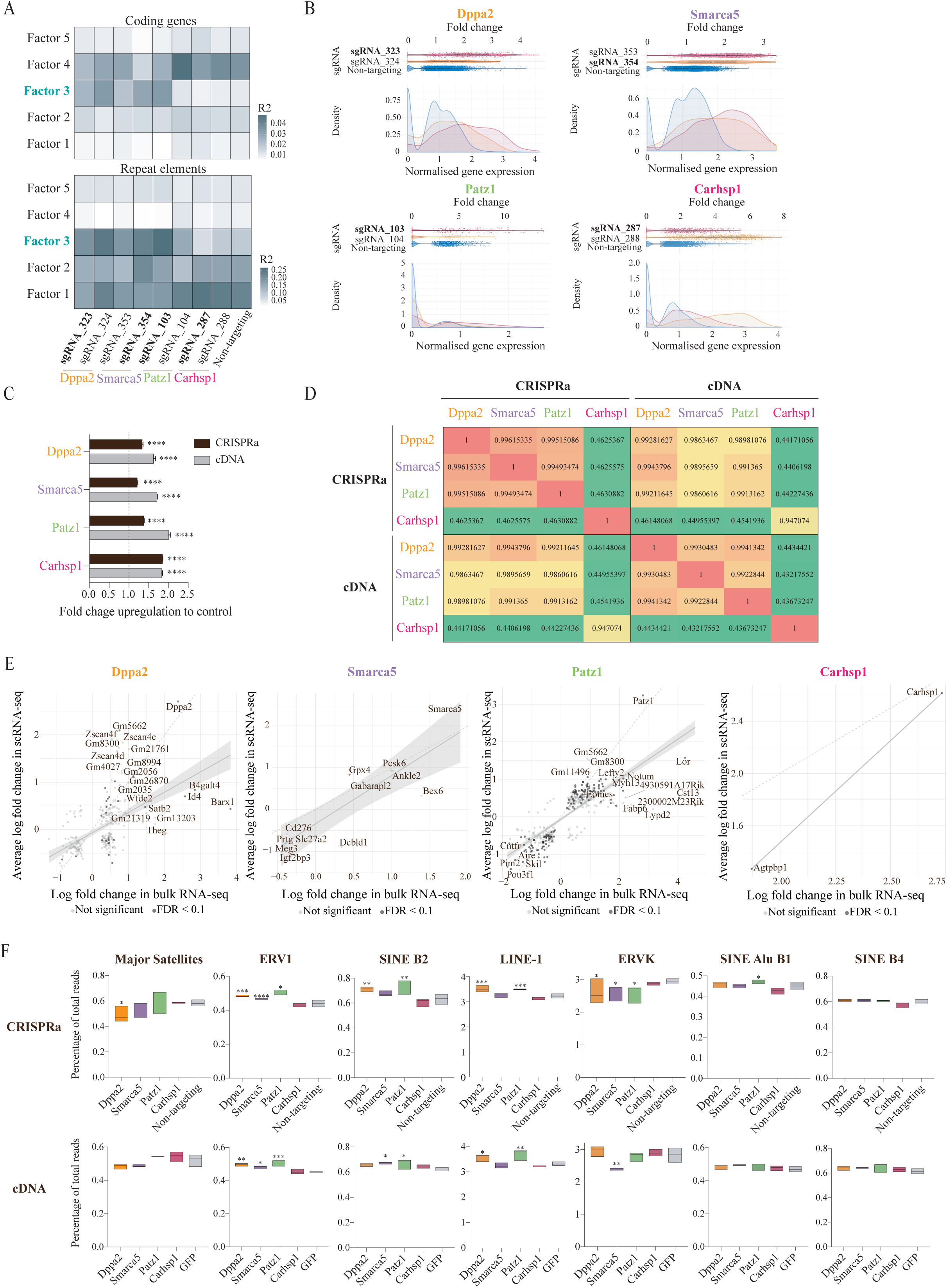
**A)** Fraction of expression variance explained (R^2^) by individual MOFA factors 1-5 for each data modality (coding genes – upper panel, and repeat elements – lower panel) in targeting sgRNAs for Dppa2, Smarca5, Patz1 and Carhsp1 and non-targeting sgRNA controls (see Table S1 for sgRNA identifiers). The sgRNA of each target gene used for validation bulk RNA-seq experiments is highlighted in bold. **B)** Dot-plot (upper panels) and density representation (lower panels) of expression levels of Dppa2 (top left), Smarca5 (top right), Patz1 (bottom left) and Carhsp1 (bottom right) in cells expressing their respective targeting sgRNAs (red and orange, see Table S1 for sgRNA identifiers) and in cells expressing non-targeting sgRNA controls (blue). In the dot-plots, each dot represents a cell plotted as a function of fold change expression to the average expression in the non-targeting sgRNA control cells. In the density plots, normalised expression is calculated as log-transformed counts per million and plotted as a function of density. The sgRNA of each target gene used for validation bulk RNA-seq experiments is highlighted in bold. **C)** Fold change upregulation of Dppa2, Smarca5, Patz1 and Carhsp1 by CRISPRa (black) and cDNA overexpression (grey) compared to controls (two non-targeting sgRNAs for CRISPRa- see Table S1- and GFP^+^-only for cDNA overexpression), measured by bulk RNA-sequencing. Data is shown as mean plus standard deviation of three biological replicates. Differences to controls are statistically significant (homoscedastic two-tailed t-test, ****: p-value < 0.0001). The sgRNAs used for CRISPRa in these experiments are described in Table S1. **D)** Heatmap of Pearson correlation coefficients between bulk gene expression profiles of Dppa2, Smarca5, Patz1 and Carhsp1 CRISPRa and cDNA overexpression. The sgRNAs used for CRISPRa in these experiments are described in Table S1. **E)** Scatterplot of log fold change values of differentially expressed genes estimated based on bulk CRISPRa RNA-sequencing data (FDR<0.1) (x axis) versus log fold change estimates for the corresponding genes in cells expressing the same sgRNA based on the CRISPRa scRNA-seq data (y axis) for the target genes Dppa2, Smarca5, Patz1 and Carhsp1. Genes that were also differentially expressed in scRNA-seq data (FDR<0.1) are labelled in dark grey whereas those genes differentially expressed in bulk RNA-sequencing data but not in scRNA-seq are labelled in light grey (not significant). For each target gene, a regression line was fitted to highlight the trend. Dashed lines mark *y = x* line. The sgRNAs used for CRISPRa in bulk RNA-sequencing experiments are described in Table S1. **F)** Box-whisker plots showing expression of different repeat families in percentage of total reads measured by bulk RNA-sequencing after CRISPRa (top panels) and cDNA overexpression (bottom panels) of Dppa2 (orange), Smarca5 (purple), Patz1 (green) and Carhsp1 (pink) and in controls (two non-targeting sgRNAs for CRISPRa and GFP^+^-only for cDNA overexpression). Statistically significant differences to controls are reported as ****: p-value < 0.0001, ***: p-value <0.001, **: p-value <0.01, * p-value <0.5, absence of stars means a non-significant difference to controls; homoscedastic two-tailed t-test. The sgRNAs used for CRISPRa in these experiments are described in Table S1.

**Supplemental Figure 6. Related to Figure 5.**
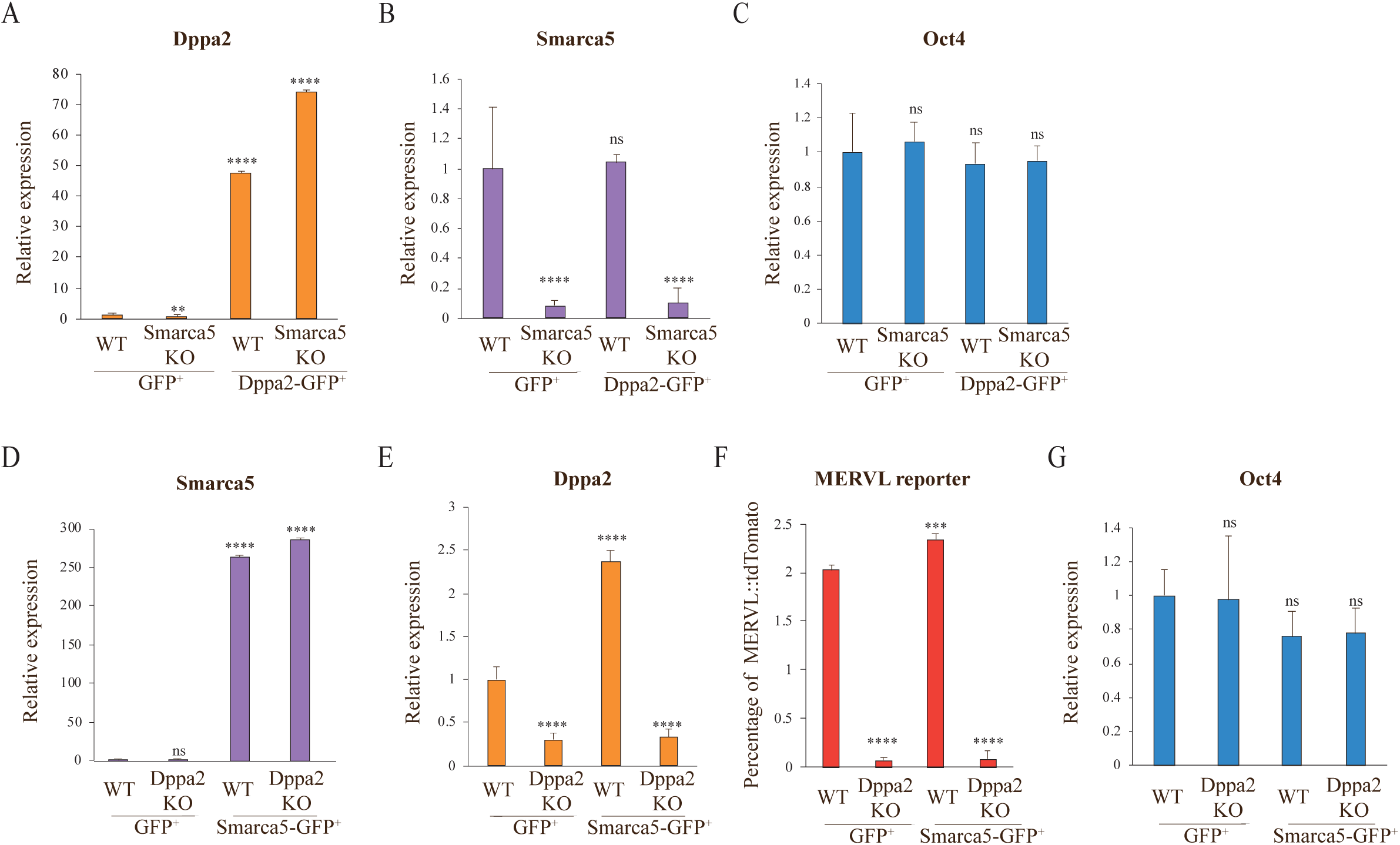
**A, B, C)** Analysis of Dppa2 (A), Smarca5 (B) and Oct4 (C) relative expression levels by quantitative reverse transcription PCR in wild-type (WT) and Smarca5 knock-out (KO) mouse ESCs after 48 hours transient transfection of GFP or Dppa2-GFP and FACS-sorting for GFP^+^ cells. **D, E, G)** Analysis of Smarca5 (D), Dppa2 (E) and Oct4 (G) relative expression levels by quantitative reverse transcription PCR in WT and Dppa2 KO mouse ESCs after 48 hours transient transfection of GFP or Smarca5-GFP and FACS-sorting for GFP^+^ cells. **F)** Percentage of GFP^+^ cells expressing a MERVL::tdTomato reporter in WT and Dppa2 KO mouse ESCs after 48 hours transient transfection of GFP or Smarca5-GFP, analysed by flow cytometry. In all panels, relative expression levels are normalised to WT cells transfected with GFP and sorted for GFP^+^; data is shown as mean plus standard deviation of three biological replicates; statistically significant differences to WT GFP^+^ control are reported (homoscedastic two-tailed t-test, **: p-value <0.01, ***: p-value <0.001, ****: p-value < 0.0001, ns: non-significant).

